# Ancient genomes redate the extinction of *Sussemionus*, a subgenus of *Equus*, to late Holocene

**DOI:** 10.1101/2021.09.13.460072

**Authors:** Dawei Cai, Siqi Zhu, Mian Gong, Naifan Zhang, Jia Wen, Qiyao Liang, Weilu Sun, Xinyue Shao, Yaqi Guo, Yudong Cai, Zhuqing Zheng, Wei Zhang, Songmei Hu, Xiaoyang Wang, He Tian, Youqian Li, Wei Liu, Miaomiao Yang, Jian Yang, Duo Wu, Ludovic Orlando, Yu Jiang

## Abstract

The exceptionally-rich fossil record available for the equid family has provided textbook examples of macroevolutionary changes. Horses, asses and zebras represent three extant subgenus of *Equus* lineage, while the *Sussemionus* subgenus is another remarkable *Equus* lineage ranging from North America to Ethiopia in Pleistocene. We sequenced 26 archaeological specimens from northern China in Holocene showing morphological features reminiscent of *Equus ovodovi*, a species representative of *Sussemionus*, and further confirmed them as this species by genetic analyses. Thus, we present the first high-quality complete genome of the *Sussemionus* that we sequenced to 12.0× depth-of-coverage and demonstrate that it survived until ∼3,500 years ago, despite the continued demographic collapse during the Last Glacial Maximum and the great human expansion in East Asia. We also confirmed the *Equus* phylogenetic tree, and found *Sussemionus* diverged from the ancestor of non-caballine equids ∼2.3-2.7 Million years ago and admixture events could have taken place between them. Our works suggest the small genetic diversity but not the enhanced inbreeding mainly limited the chances of survival of the species, and illustrates how ancient DNA can inform on extinction dynamics and the long-term resilience of species surviving in cryptic population pockets.

## Introduction

Today, all of the seven extant species forming the horse family belong to one single genus, *Equus*. It emerged in North America some 4.0-4.5 million years ago (L. Orlando et al., 2013), and first spread to Eurasia ∼2.6 million years ago, via the Beringia land bridge (Lindsay, Opdyke, & Johnson, 1980). This first vicariance and expansion out-of-America led to the emergence of the ancestors of zebras, hemiones and donkeys, a group collectively known as non-caballine equids. Another expansion through Beringia occurred later, bringing caballine equids (i.e. those most closely related to the horse) into the Old-World, where they survived until their domestication some ∼5,500 years ago (Gaunitz et al., 2018; Outram et al., 2009).

In the recent years, ancient DNA (aDNA) data have revealed that the genetic diversity of non-caballine *Equus* was considerably larger diversity in the past than it is today (Librado & Orlando, 2020; Ludovic Orlando, 2020), especially as the first mitochondrial DNA (mtDNA) data of *Equus* (*Sussemionus*) specimens were collected (hereafter referred to as Sussemiones) (Eisenmann, 2010). This lineage radiated across North America, Africa, and Siberia, and developed multiple adaptations to a whole range of arid and humid environments (Eisenmann, 2010). Sussemiones was first believed to have become extinct during the Middle Pleistocene as the last known specimen showing typical morpho-anatomy dated back to approximately 500,000 years ago (Vasiliev, 2013). However, DNA results obtained on multiple osseous remains within the radiocarbon range and showing morphological traits reminiscent of the Eurasian Sussemiones species indicated that the lineage in fact survived until the Late Pleistocene (Druzhkova et al., 2017; L. Orlando et al., 2009; Vilstrup et al., 2013; Yuan et al., 2019). Pioneering publications indicated survival dates 40-50 kya in southeastern Siberia, Russia (Proskuryakova cave) (L. Orlando et al., 2009; Vilstrup et al., 2013), ∼32 kya at the Denisova cave (Druzhkova et al., 2017), and ∼12.6 kya at northeastern China (Yuan et al., 2019).

Despite an abundant fossil material, only a limited number of Sussemiones specimens have been investigated for Ancient mitochondrial DNA (aDNA), which showed that Sussemiones formed a non-caballine equine lineage. However, the exact placement of Sussemiones could not be resolved (Heintzman et al., 2017; L. Orlando et al., 2009; Vilstrup et al., 2013). In this study, we have carried out archaeological excavations in three Holocene sites in China, and uncovered equine samples showing distinct morphological features when compared to horses and donkeys (Figure 1—figure supplement 1). The whole mitochondrial and nuclear genome data allowed us to unveil the phylogenetic placement of Sussemiones within the *Equus* evolutionary tree, the timing of its divergence to other non-caballine equids, signatures of demographic collapse and adaptations specific to this lineage, and the extinction dynamics of this lineage.

## Results

### Archaeological samples and sequencing data

All the equine specimens investigated in this study were excavated from three archaeological sites in China (Figure 1 and Table S1) (Honghe, Heilongjiang Province (Figure 1—figure supplement 2); Muzhuzhuliang, Shaanxi Province; Shatangbeiyuan, Ningxia Province). They showed morphological and genetic signatures distinct from those of extant horses and donkeys. The morphological differences were especially marked in the second and third molars, which appeared to be smaller than in modern horses, and were reminiscent of the third molars paracones and metacones observed in Sussemiones specimens (Figure 1B). Combined, these samples were radiocarbon dated to 3,477-4,481 calibrated years before the present (cal BP), including the latest sample HH13H with 3,477-3,637 cal BP (Table S2). They could, thus, represent some of the latest surviving Sussemiones individuals prior to their extinction.

**Figure 1.**
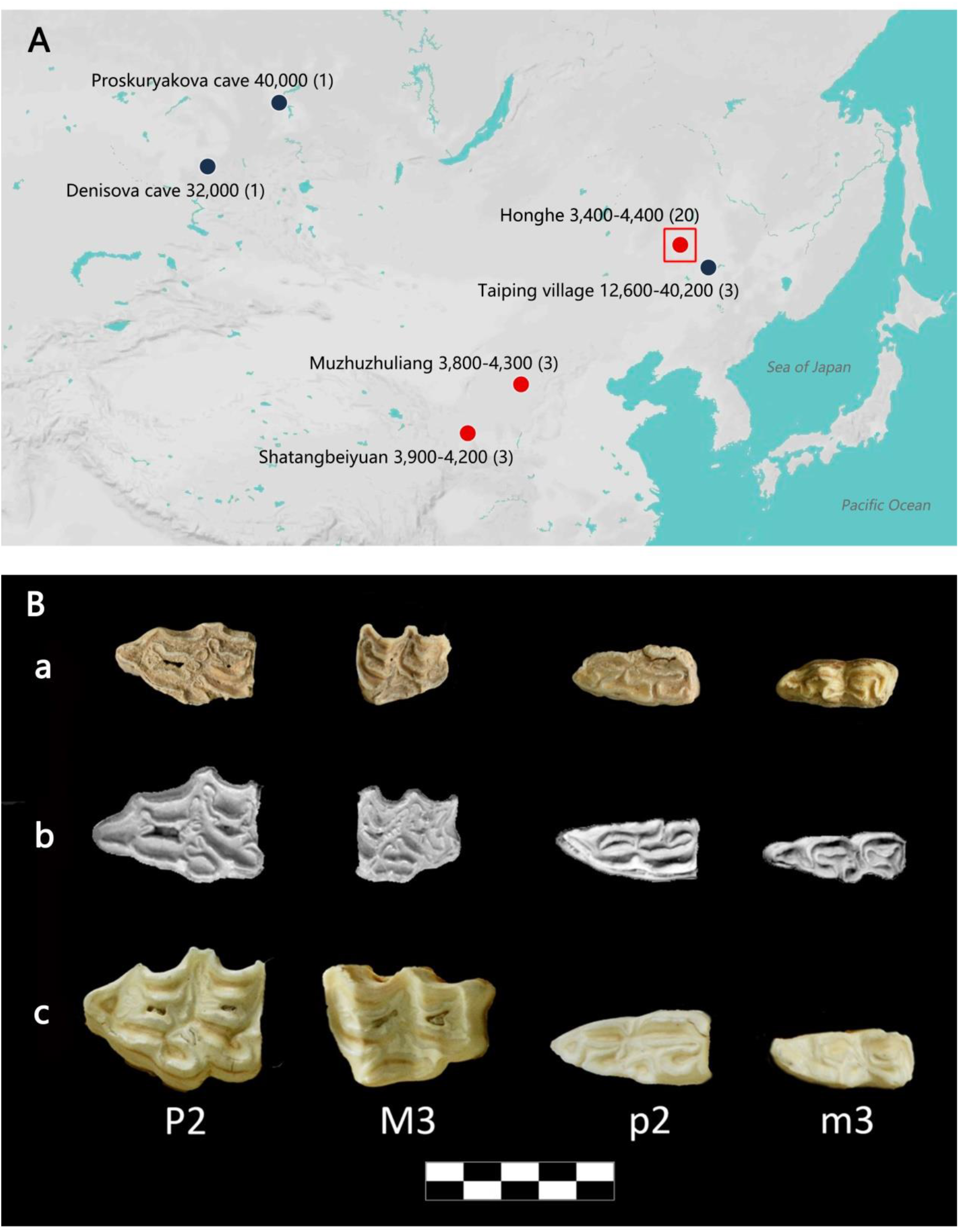
Sampling distribution and archaeological research of *E.* (*Sussemionus*) *ovodovi*. (**A**) *E.* (*Sussemionus*) *ovodovi* geographic range. The three red circles indicate the archaeological sites analyzed in this study. The site (Honghe) that delivered the complete genome sequence at 12.0-fold average depth-of-coverage (HH06D) is highlighted with a square. The black circles indicate sites that provided complete mitochondrial genome sequences in previous studies (Druzhkova et al., 2017; L. Orlando et al., 2009; Vilstrup et al., 2013; Yuan et al., 2019). The temporal range covered by the different samples analyzed is given in years before present (YBP) and follows the name of each site. Numbers between parentheses indicate the number of samples for which DNA sequence data could be generated. (**B**) *Facies masticatoria dentis* of P2, M3, p2 and m3 for the *E. (Susseminous) ovodovi* samples of the Honghe site analyzed here (a), *E. Sussemionus* (b), and *E. caballus* (c). Teeth from the right side are shown, except for *E. Sussemionus*. The erupted teeth of the samples of the Honghe site appear to be smaller than those of the *E. Sussemionus* specimen. The following figure supplements are available for figure 1: **Figure supplement 1.** Archaeological material investigated in this study. **Figure supplement 2.** Aerial view of the Honghe site.

We next aimed at genetically characterizing and identifying the taxonomic status of these specimens using high-throughput DNA sequencing technologies. We extracted ancient DNA from a total of 26 specimens and sequenced the whole nuclear genome at ∼0.002 to 12.0 times coverage, including three samples from Honghe provided 12.0×, 3.5× and 1.0× nuclear genome (Table S1). Comparison of the X chromosome and autosomal coverage revealed the presence of 15 males and 11 females.

### Taxonomic status

To assess whether the sequenced specimens belonged to the same taxonomic group or comprised different species, we carried out a Principle Component Analysis (PCA) including all the equine species sequenced at the genome level (depth of coverage ≥ 1×) (Figure 2A and Figure 2—figure supplements 3 and 4). For this, we downloaded 11 previously-published equine genomes representing all extant species of equids and the extinct quagga zebra (Huang et al., 2015; Jonsson et al., 2014; Kalbfleisch et al., 2018; L. Orlando et al., 2013; Renaud et al., 2018) (Table S3). All the Chinese specimens analyzed in this study were found to cluster together along the first two PCA components, in a group that was distinct from all other equine species (Figure 2A and Figure 2—figure supplement 3) but closer to non-caballine equine species than to the horse (Figure 2A). This suggested that they were all members of a unique taxonomic group, most related to non-caballine equids.

**Figure 2.**
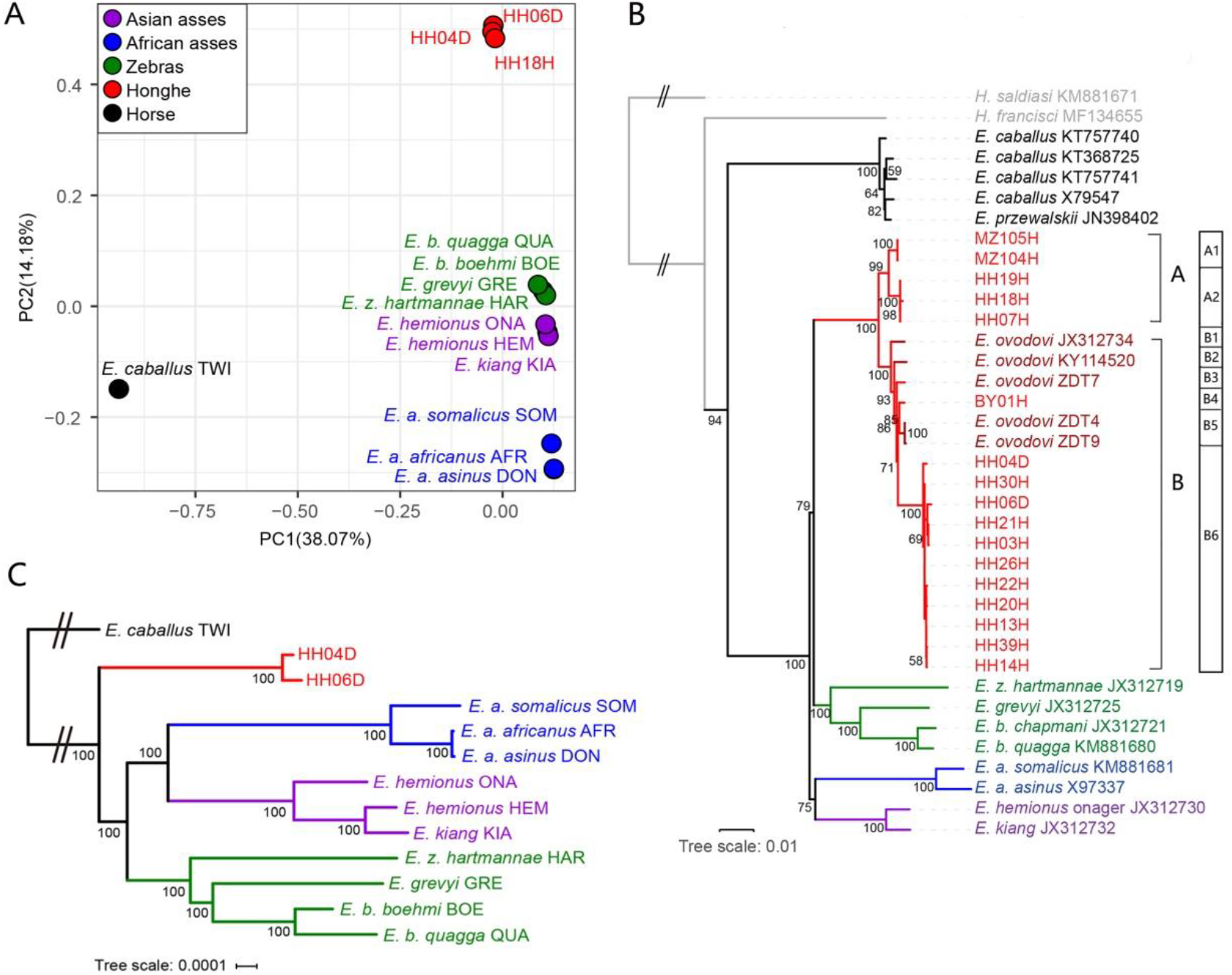
Genetic relationships within the *Equus* genus. Honghe (HH), Muzhuzhuliang (MZ) and Shatangbeiyuan (BY) specimens are shown in red, while Asian asses, African asses, zebras and horses are shown in whereas purple, blue, green and black, respectively. (**A**) PCA based on genotype likelihoods, including horses and all other extant non-caballine lineages (16,293,825 bp, excluding transitions). Only specimens whose genomes were sequenced at least to 1.0× average depth-of-coverage are included. (**B**) Maximum likelihood tree based on 5 mitochondrial partitions (representing a total of 15,399 bp). Previously published *E. ovodovi* sequences are shown in deep red. The tree was rooted using *Hippidion saldiasi* and *Haringtonhippus francisci* as outgroups (not shown). Node supports were estimated from 1,000 bootstrap pseudo-replicates and are displayed only if greater than 50%. The black line indicates the mitochondrial haplogroup A and B. (**C**) Maximum likelihood tree based on sequences of 19,650 protein-coding genes with specimens sequenced at least 3.0× average coverage (representing 32,756,854 bp). The following figure supplements are available for figure 2: **Figure supplement 1.** DNA Damage patterns for HH06D. **Figure supplement 2.** Error profiles of the 26 ancient genomes characterized in this study. **Figure supplement 3.** Principal Component Analysis (PCA) based on genotype likelihoods using the horse reference genome. **Figure supplement 4.** Principal Component Analysis (PCA) based on genotype likelihoods using the donkey reference genome. **Figure supplement 5.** RAxML-NG (GTR+GAMMA model) Maximum Likelihood phylogeny of complete mitochondrial sequence data. **Figure supplement 6.** Bayesian mitochondrial phylogeny based on 6 partitions and using *Hippidion Saldiasi* as outgroup. **Figure supplement 7.** Exome-based Maximum likelihood phylogeny rooted by the horse lineage. **Figure supplement 8.** Treemix analysis of based on genome-wide SNP data conditioned on transversions using the horse reference genome. **Figure supplement 9.** Treemix analysis of based on genome-wide SNP data conditioned on transversions using the donkey reference genome.

Maximum likelihood (ML) phylogenetic analyses including the nearly-complete 17 mitochondrial genomes reported in this study (Table S1, depth of coverage no less than 1×) confirmed their clustering with non-caballine equids, within a single monophyletic group that also included five previously-characterized Sussemiones specimens (Figure 2B and Figure 2—figure supplements 5 and 6). This grouping was supported with maximal (100%) bootstrap support. This, and the PCA clustering, indicated that the different excavation sites investigated in this study in fact all provided specimens that belonged to the *E.* (*Sussemionus*) *ovodovi* species.

We also inferred maternal lineages of *E. ovodovi* using complete mitogenomes. Phylogenetic analyses showed that the species were mainly divided into two major haplogroups (A and B), of which most of the specimens were characterised by the haplogroup B, including all previously reported sequences, whereas 2 Muzhuzhuliang and 3 Honghe individuals belonged to the new haplogroup A (Figure 2B). Further distinction could also be carried out in the two haplogroups (Figure 2B).

To further assess phylogenetic affinities, we used the two genomes characterized to at least 3× average depth-of-coverage (HH04D and HH06D) to replace Sussemiones within the equine phylogenetic tree. To achieve this, we used ML phylogenetic reconstruction and an alignment of the coding sequences of the protein coding genes ( Figure 2C and Figure 2—figure supplement 7A). This showed that the Chinese ancient specimens branched off before the radiation leading to modern asses and zebras (Figure 2C). Similar tree topologies were recovered using whole-genome SNPs by TreeMix (Pickrell & Pritchard, 2012) (Figure 2—figure supplements 8 and 9). Combined with the analysis of the occlusal surface of the molars, in particular the absence of the caballine notch, the shape of metacones and protocones, and the reduced tooth size (Figure 1B), our analyses allowed us to conclude that the material analyzed represented small specimens of the extinct *Equus* (*Sussemionus*) *ovodovi*. This lineage, thus, survived in China during the Holocene, and until cal 3,477-4,481 cal BP, which is approximately ∼8,300-9,300 years after the latest known specimen to date (Druzhkova et al., 2017; L. Orlando et al., 2009; Vilstrup et al., 2013; Yuan et al., 2019).

### Interspecies admixture and demographic modeling

Bifurcating trees fail to capture possible admixture events between lineages. Yet, previous research has unveiled pervasive admixture within equids, even amongst extant equids showing different chromosomal numbers (Jonsson et al., 2014). We thus next assessed whether the genomic data showed evidence for gene flow between Sussemiones and other non-caballine equids. To achieve this, we first applied D-statistics (Soraggi, Wiuf, & Albrechtsen, 2018) to the genome sequence underlying 26 individual genomes and detected that *E. ovodovi* shared an excess of derived polymorphisms with asses than relative to zebras (Figure 3—figure supplements 1 and 2). This suggested that at least one admixture event could have taken place between Sussemiones and the ancestor of asses after their divergence from zebras.

We next leveraged the ancient genome characterized to high depth-of-coverage (HH06D) to reconstruct the equine demographic history using G-PhoCS (Gronau, Hubisz, Gulko, Danko, & Siepel, 2011). More specifically, we first selected members of each equine lineage representing a total number of 10 genomes, and assumed that the genus *Equus* emerged some 4.0-4.5 Mya, following previous estimates (L. Orlando et al., 2013). G-PhoCS analysis confirmed previous analyses indicating that the zebras and asses linages diverged ∼2.0 Mya and that the deepest divergence within zebras and asses took place prior to ∼1.5 Mya (Jonsson et al., 2014) (Figure 3). It revealed that the Sussemiones lineage diverged from the ancestor of extant non-caballine equids ∼2.3-2.7 Mya, in line with the fossil record (Eisenmann, 2010). Allowing for migrations provided support for gene flow between Sussemiones and the ancestor of asses and zebras (Figure 3). However, weak to no migrations were detected between Sussemiones and extant equids (Table S6). Importantly, the admixture between Sussemiones and the ancestor of asses seems to have been stronger than that between Sussemiones and the ancestor of zebras, in line with the results of D-statistics. G-PhoCS also supported the presence of significant unidirectional gene-flow prior to ∼2.3-2.7 Mya, from the horse branch into the ancestral branch to all non-caballine equids, including Sussemiones (total migration rate 2.2-9.2%, Table S7). This is consistent with previous HMMCoal analyses applied to whole genome sequences of all extant equine species, which indicated significant gene-flow between the deepest branches of the *Equus* phylogenetic tree until 3.4 Mya, mostly from a caballine lineage into the ancestor of all non-caballine equids (Jonsson et al., 2014).

**Figure 3.**
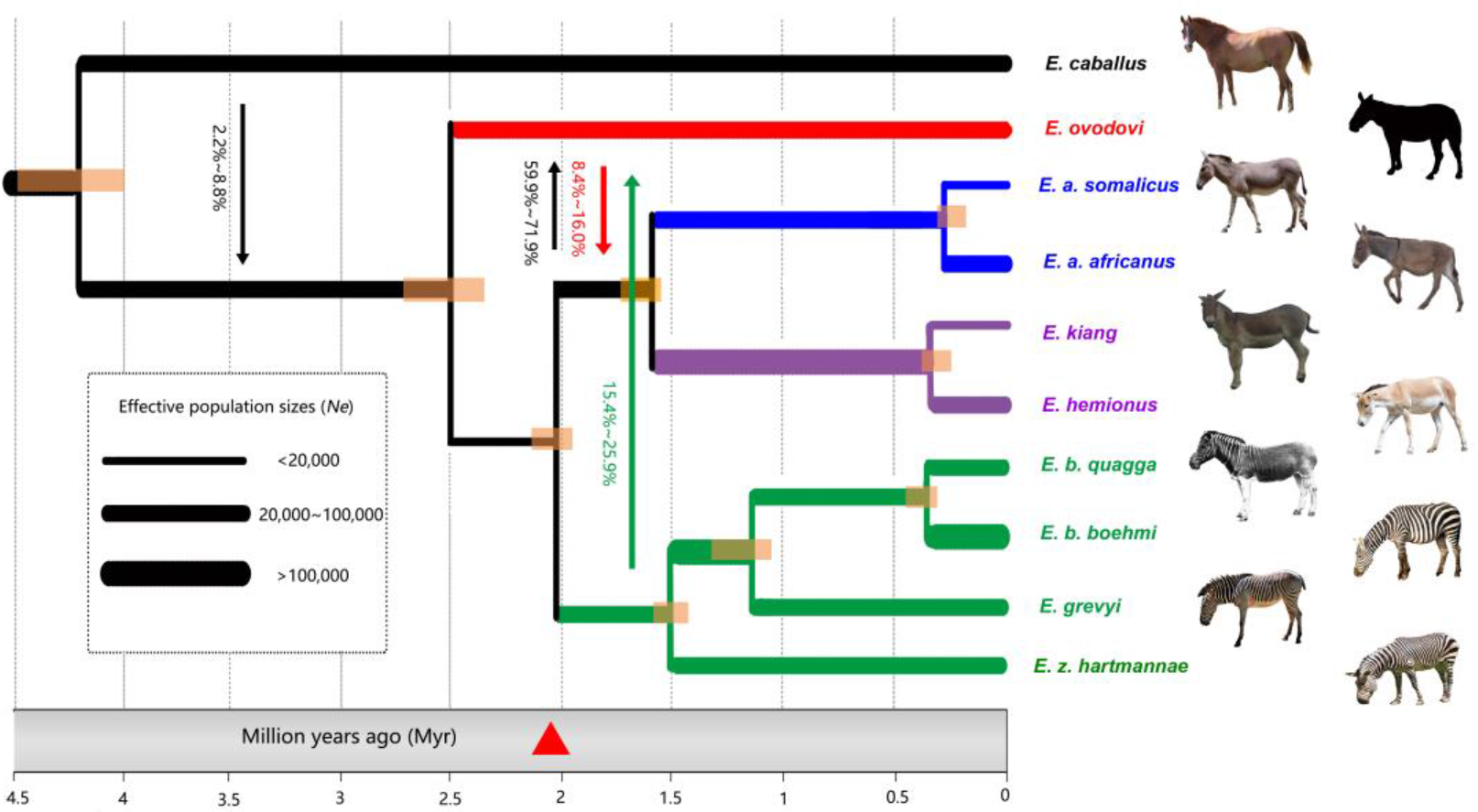
Demographic model for extinct and extant equine lineages as inferred by G-PhoCS (Gronau et al., 2011). Node bars represent 95% confidence intervals. The width of each branch is scaled with respect to effective population sizes (*N_e_*). Independent *N_e_* values were estimated for each individual branch of the tree, assuming constant effective sizes through time. Migration bands and probabilities of migration (transformed from total migration rates) are indicated with solid. The red triangle indicates the earliest *Sussemionus* evidence found in the fossil record. (Images: *E. caballus* by Infomastern, *E. a. somalicus* by cuatrok77, *E. kiang* by Dunnock_D, *E. a. africanus* by jay galvin, *E. hemionus* by Cloudtail the Snow Leopard, *E. z. hartmannae* by calestyo, *E. b. quagga* by Internet Archive Book Images, *E. b. boehmi* by GRIDArendal, and *E. grevyi* by 5of7.) The following figure supplements are available for figure 3: **Figure supplement 1.** *D-*statistics in the form of (zebra, ass; *E. ovodovi*, outgroup), using sequence alignments against the horse reference genome. **Figure supplement 2.** *D-*statistics in the form of (zebra, ass; *E. ovodovi*, outgroup), using sequence alignments against the donkey reference genome. **Figure supplement 3.** NJ tree of selected samples based on 15,324 candidate ‘neutral’ loci identified using sequence alignments against the horse reference genome.

### Dynamic demographic profiles, heterozygosity and inbreeding levels

We next leveraged the high-coverage Sussemiones genome characterized here to explore further the demographic dynamics of this lineage until its extinction. When modeled as constant through time, population sizes in G-PhoCS indicated that most lineages, including Sussemiones, consisted of small populations, excepting the Burchell’s zebra (Table S8). Pairwise Sequential Markovian Coalescent (PSMC) analyses, however, provided us evidence for population size variation through time. First, the Sussemiones demographic trajectory was found to diverge from that of other non-caballine equids (specifically, *E. hemionus*) after ∼2.0 Mya, confirming the divergence date estimate retrieved by G-PhoCS (Table S8). Second, we found that the Sussemiones demographic trajectory constantly increased during the last million year but stay at a level which was lower than that of other lineages for a long time, until it reached a peak between 74-84 kya. It was, then, followed by an approximately 45-fold collapse until 13 kya (Figure 4). The Sussemiones population size experienced a 1.86-fold collapse between 35-42 kya, which is almost coincident with the timing of the great human expansion to Eurasia (ie. 35-45 kya, (Henn, Cavalli-Sforza, & Feldman, 2012)). The lineage maintained extremely reduced population sizes through the Last Glacial Maximal (LGM, 19-26 kya) (Clark et al., 2009) and the Holocene, until it finally became extinct.

**Figure 4.**
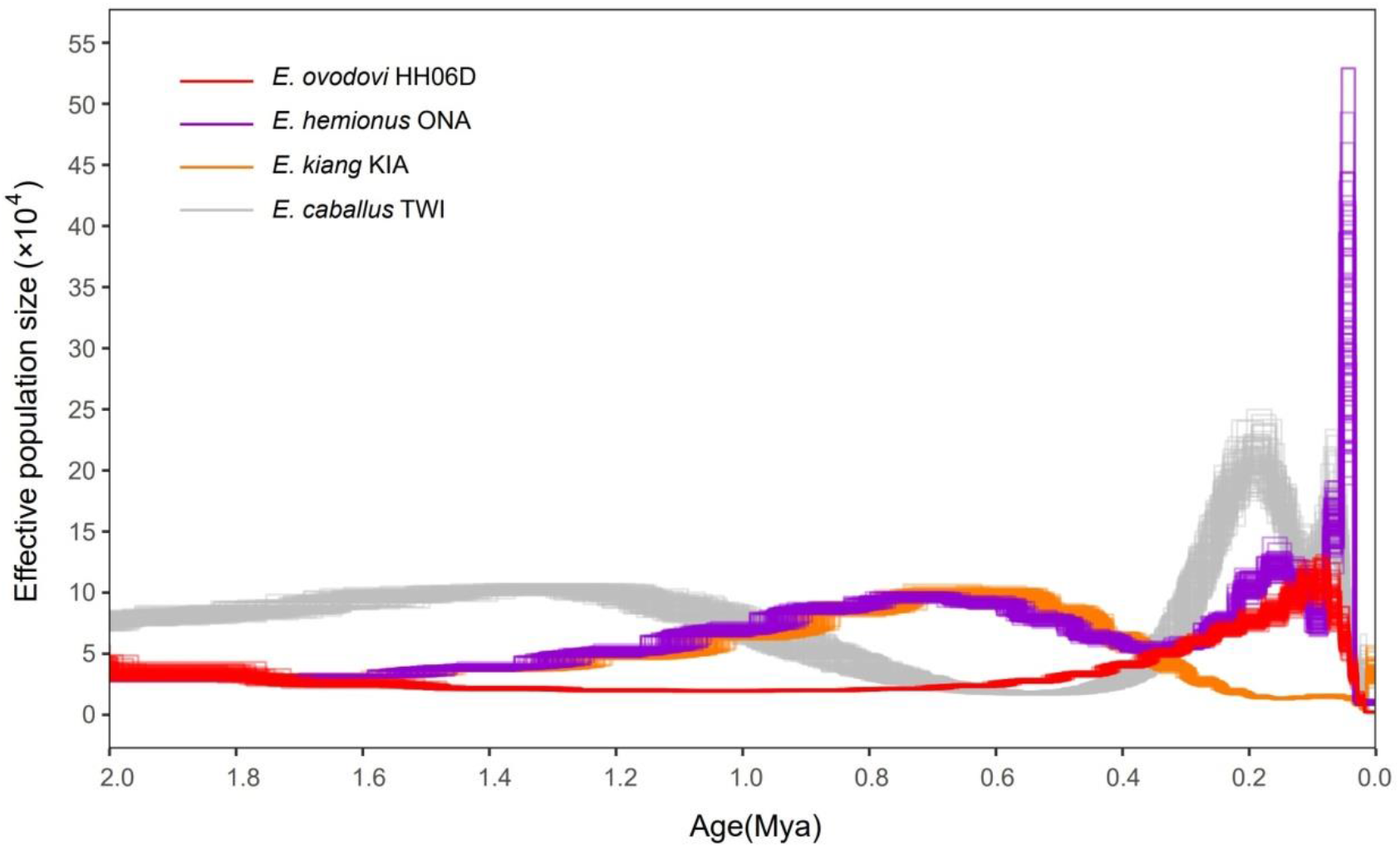
PSMC profiles (100 bootstrap pseudo-replicates) of four Eurasian equine species (*E. ovodovi* HH06D, *E. caballus* TWI (Kalbfleisch et al., 2018), *E. hemionus* ONA and *E. kiang* KIA) (Jonsson et al., 2014). The y axis represents effective population size (×10,000), and the x axis represents millions of years before present. Faded lines show bootstrap values. The following figure supplements are available for figure 4: **Figure supplement 1.** PSMC bootstrap pseudo-replicates for samples with and without transitions. **Figure supplement 2.** Determining the uniform false-negative rate (uFNR) that was necessary for PSMC scaling.

Importantly, the sample sequenced to sufficient coverage (HH06D) showed minimal heterozygosity and moderate inbreeding levels identified by the fraction of the segments within ROH (Figure 5). Strikingly, this is true in spite of the increased sequencing error rates of this genome, which likely inflate our estimates. The limited population sizes and genetic diversity but not the enhanced inbreeding may have limited the chances of survival of the species, ultimately leading to extinction.

**Figure 5.**
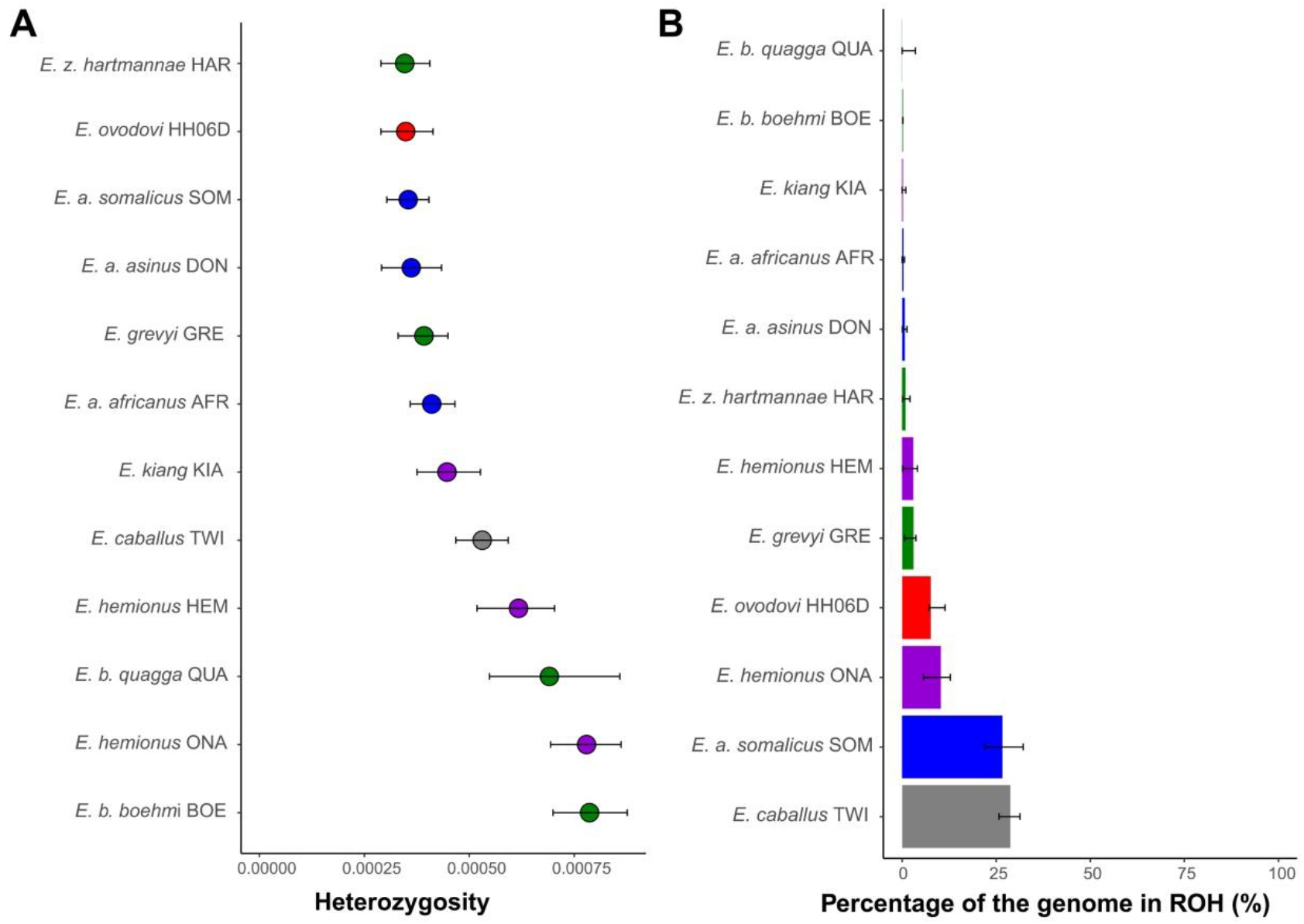
The Heterozygosity and Inbreeding levels of extinct and extant equine lineages. (**A**) Individual heterozygosity outside Runs-of-Homozygosity (ROH). (**B**) Fraction of the genome in ROH. Estimates were obtained excluding transitions and are shown together with their 95% confidence intervals. The colors mirror those from Figure 2. The following figure supplements are available for figure 5: **Figure supplement 1.** Heterozygosity rates outside Runs-of-Homozygosity (ROH) together with 95% confidence intervals. **Figure supplement 2.** The fraction of the genome segments consisting of ROHs together with 95% confidence intervals.

## Discussion

### Phylogenetic placement of *Equus* (*Sussemionus*) *ovodovi*

In this study, we have characterized the first nuclear genomes of the now-extinct equine lineage, *E.* (*Sussemionus*) *ovodovi*, the last surviving member of the subgenus *Equus Sussemionus*. We demonstrated that this lineage survived in China well into the Holocene with the most recent specimens analyzed dating to ∼3,477-3,637 cal BP. This is almost 9,000 years after the latest specimens previously documented in the fossil record (Druzhkova et al., 2017; Vilstrup et al., 2013; Yuan et al., 2019). Our work, thus, shows that *Sussemionus* represents the last currently known *Equus* subgenus to become extinct. Our work also adds to the list of recently identified members of the horse family that were still alive at the time horses and donkeys were first domesticated, some ∼5,500 years ago (Fages et al., 2019; Gaunitz et al., 2018; Rossel et al., 2008). In contrast to those divergent members that were identified in Siberia and Iberia and both belonged to the horse species (Fages et al., 2019; Schubert, Jonsson, et al., 2014), Sussemiones members were most closely related to non-caballine equids. This is in agreement with previous studies (Der Sarkissian et al., 2015; Druzhkova et al., 2017; Heintzman et al., 2017; L. Orlando et al., 2009; Vilstrup et al., 2013; Yuan et al., 2019), which could, however, not fully resolve the exact phylogenetic placement of this species within non-caballine clade lineages as topological tests based on mitochondrial genomes received low confidence support (Der Sarkissian et al., 2015; Druzhkova et al., 2017; Heintzman et al., 2017; L. Orlando et al., 2009; Vilstrup et al., 2013; Yuan et al., 2019). Our study solved this question by reporting the first whole genome phylogeny of Sussemiones, which confirmed with maximal bootstrap support this species as a unique basal lineage of non-caballine equids.

### Suitable habitat and geographic distribution

Previous zooarchaeological and environmental research indicated an ecological range for Sussemiones overlapping with the grasslands located east of the Altay Mountains and west of the Yenisei River during the Late Pleistocene (Khenzykhenova et al., 2016; Malikov, 2016; Plasteeva, 2015; Shchetnikov, Klementiev, Filinov, & Semeney, 2015; Shunkov, 2018). Recent research also reported this species in northeastern China (∼12,600-40,200 YBP), where similar climatic and ecological conditions were found at the time (Yuan et al., 2019). It could, thus, be speculated that Sussemiones was adapted to an environment with moderately dry climatic conditions and steppe landscapes (Yuan et al., 2019). However, our study identified Sussemiones in three late Holocene sites from China that have mild and humid environmental conditions. In addition, two distinct mitochondrial haplogroups from 22 individuals have been defined from the six known sites, suggesting that Sussemiones had adapted to different environmental regions. It also suggests that the species could adapt to a wider variety of habitats than previously hypothesized, and rejects the contention that the species became extinct as it could not survive in warmer climatic conditions (Yuan et al., 2019).

Interestingly, the Sussemiones specimens identified in this study were excavated from sites in northeastern China located at almost the same latitude as those Sussemiones localities known so far from Russia, but also at lower latitudes (Figure 1A). This implies that the geographic range of *E. ovodovi* was larger than previously expected and included at least Northern China and Southern Siberia. In the absence of identified fossils from Mongolia, whether those two regions were in contact or separated remains unknown. Further work is necessary to establish whether or not the species survived in other pockets both within and outside China.

### Demographic history with ancestral interspecific admixture

Our analyses reveal that the divergence between Sussemiones and the most recent common ancestor of all extant non-caballine equids took place some ∼2.3-2.7 Mya, right before the divergence of zebras and asses. Post-divergence admixture events with the lineage ancestral to asses and zebras on the one hand, and the lineage ancestral to all extant zebras, were also identified (Figure 3 and Table S7). Our results, thus, reveal non-caballine ancestral lineages occupying partly sympatric distributions that were, consequently, different than those of their descendants, in which zebras are restricted in Africa and Asian asses in Asia. Whether the admixture events identified here directly involved the Sussemiones lineage or one (or more) ghost lineage(s) closely related to Sussemiones requires further research.

### Limited genetic diversity before extinction

The demographic profile of Sussemiones shows that after the peak of population size culminating some ∼74 kya, Sussemiones went through a slow and continuous decline until 13 kya (Figure 4). This time period encompasses several major climate changes (especially the LGM) and the great human expansion to Eurasia (∼35-45 kyr BP) (Henn et al., 2012). The effective size of Sussemiones populations that survived in Northern China until at least ∼3,500 years ago, remained extremely small, as indicated by their extremely reduced heterozygosity levels compared to other extant and extinct equine species, although extensive inbreeding was not detected (Figure 5). So combined with a degree of inbreeding, the reduced genetic diversity available ultimately resulted in the extinction of the lineage, in a process reminiscent of what was previously described for the woolly mammoth (Palkopoulou et al., 2015).

In conclusion, our study clarifies the phylogenetic placement, speciation timing and evolutionary history of the now-extinct *Equus Sussemionus* equine subgenus. This group did not remain in reproductive isolation from other equine lineages, but contributed to the genetic makeup of the ancestors of present-day Asiatic asses, while receiving genetic material from the ancestors of African zebras. This supports geographic distributions at least partly overlapping at the time, thus, not identical to those observed today. The species demographic trajectory experienced a steady decline from ∼74 kya and during a period witnessing both important climatic changes and the Great human expansion across Asia (Henn et al., 2012). It survived with minimal genetic diversity the Pleistocene-Holocene transition, and for at least eight millennia before it became extinct, which providing insights into the extinction of large animals since Holocene.

## Additional information

### Acknowledgments

We thank High-Performance Computing (HPC) of Northwest A&F University (NWAFU) for providing computing resources.

### Funding

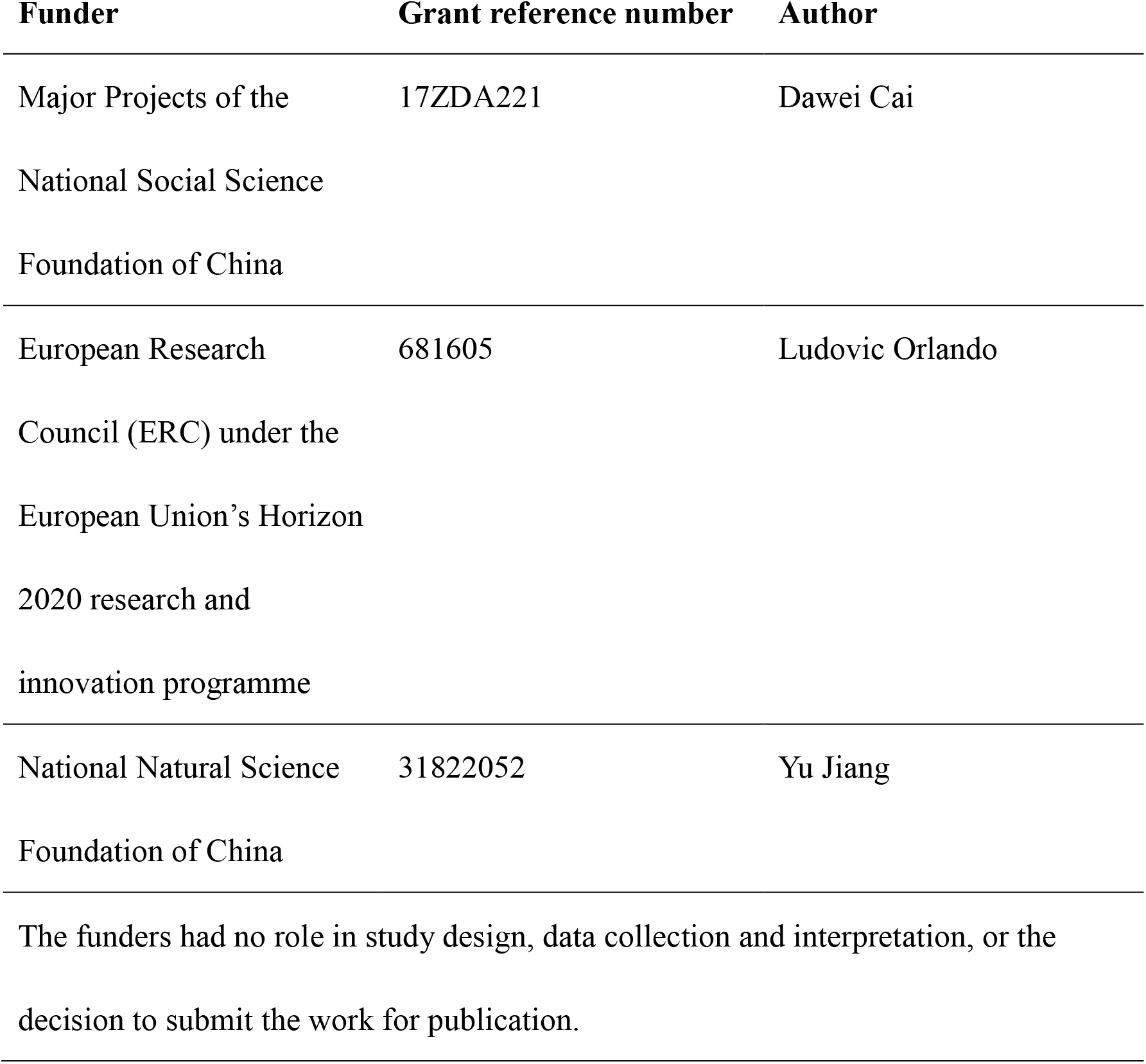

## Additional files

### Supplementary files

- Supplementary file 1. Supplementary tables that support the analysis and results above.
- Transparent reporting form

## Materials and Methods

### Genome Sequencing

Prior to DNA extraction, the outer surface of the sample was cleaned with a brush. The cleaned sample was subsequently cut into smaller pieces and soaked in 10% bleach for 20 min, rinsed with ethanol and distilled water, and then UV-irradiated for 30 min on each side. Finally, powder was obtained by drilling with a dental drill (Traus 204, Korea). Ancient DNA was extracted from the sample powder by using a modified silica spin column method (Yang, Eng, Waye, Dudar, & Saunders, 1998) in dedicated ancient DNA facilities at Jilin University (JLU). For each specimen, a total of 200mg powder was added with 3.9ml EDTA (0.465mol/L) and placed in the refrigerator at 4 ° C for 12 hours for decalcification, and then 0.1 ml Proteinase K (0.4mg/mL) were added and incubated overnight in a rotating hybridization oven at 50℃ (220rpm/min). After centrifugation, the supernatant was transferred into an Amicon® Ultra-4 centrifugal filter device (Merck Millipore Ltd, 10000 Nominal Molecular Weight Limit), reduced to less than 100ul, and purified with QIAquick® PCR Purification Kit (QIAGEN), according to the manual instructions.

Before preparation of DNA libraries, we first PCR targeted short fragments of the mitochondrial hypervariable region to select those samples positive for the presence of equine DNA (which was further confirmed through Sanger sequencing). For this, we used the oligonucleotide primers L15473 5’ –CTTCCCCTAAACGACAACAA–3’ and reverse primer H15692 5’ –TTTGACTTGGATGGGGTATG–3’ ; and forward primer L15571 5’ –AATGGCCTATGTACGTCGTG–3’ and reverse primer H15772 5’ –GGGAGGGTTGCTGATTTC–3’from (Dawei et al., 2007), and the amplification conditions therein.

Double-stranded single-indexed libraries were prepared using NEBNext® Ultra^TM^ II DNA Library Prep Kit for Illumina® (NEB #E7645S) and NEBNext® Multiplex Oligos for Illumina® Index Primers Set 1 and 2 (NEB #E7335S, #E7500S), following the manufacturer’s instructions with minor modifications. Specifically, the extracted DNA (50ul) were end repaired and A-tailed by adding 7μl of NEBNext Ultra II End Prep Reaction Buffer and 3 μl of NEBNext Ultra II End Prep Enzyme Mix, and incubated for 40 min at 20 °C and then 30 minutes at 65 °C. The adaptor was ligated to the dA-tailed DNA fragments by adding 30ul of NEBNext Ultra II Ligation Master Mix, 1ul of NEBNext Ligation Enhancer and 2.5ul of NEBNext Adaptor for Illumina (Dilution 1:10), and incubated for 20 min at 20 °C. The adaptor was then linearized by adding 3ul of USER™ Enzyme and performing an incubation for 15 min at 37°C. The adaptor-ligated DNA were cleaned without size selection using the MinElute® PCR Purification Kit (QIAGEN, Germany), following the instructions provided by the manufacturer. PCR enrichment was performed by using 30ul of NEBNext Ultra II Q5 Master Mix, 1ul of Index Primer, 1ul of Universal PCR Primer and 18ul of adaptor-ligated DNA. PCR cycling conditions comprised an initial denaturation at 98°C for 30s, 14-16 cycles of 98°C for 10s, 65°C for 75s, and a final extension at 65°C for 5min. PCR amplified DNA libraries were purified using Agencourt AMPure XP Beads, following the manufacturer’s instructions, and Illumina sequencing was performed on HiSeq X Ten platform using 150bp paired-end reads. Overall, we sequenced a total of 28 DNA libraries and generated 2,727,843,803 read pairs.

All pre-PCR procedures were conducted in a dedicated ancient DNA laboratory at JLU that is physically separated from the post-PCR laboratory. To remove potential contaminant DNA, working areas and benches were frequently cleaned with bleach and UV exposure. Lab experiments were carried out wearing full body suits, facemasks, and gloves. To detect contamination, mock blank controls were included in each experimental step, including DNA extraction, DNA library preparation and PCR setup.

### Data Processing

Sequencing reads were processed and aligned against the horse (EquCab3.0 (Kalbfleisch et al., 2018)) and donkey (Renaud et al., 2018) reference genomes using the PALEOMIX pipeline (Schubert, Ermini, et al., 2014) with default parameters, except that we followed the recommendations from (Schubert et al., 2012) and disabled seeding. Briefly, paired-end (PE) reads longer than 25 nucleotides were trimmed with AdapterRemoval v2.2 (Schubert, Lindgreen, & Orlando, 2016) and aligned against the reference genomes using BWA (Li & Durbin, 2009), retaining alignments with mapping qualities superior to 25. PCR duplicates were then removed using Picard MarkDuplicates (http://broadinstitute.github.io/picard/). Finally, all ancient and modern reads were locally realigned around indels using GATK (McKenna et al., 2010).

Postmortem DNA damage and average sequencing error rates were determined with mapDamage2.0 (Jonsson, Ginolhac, Schubert, Johnson, & Orlando, 2013) (Figure 2—figure supplement 1) and ANGSD (Korneliussen, Albrechtsen, & Nielsen, 2014) (Figure 2—figure supplement 2), respectively. Further rescaling and trimming procedures were implemented following (Gaunitz et al., 2018) to limit the impact of remnant nucleotide mis-incorporations in subsequent analyses. For each of the DNA libraries examined, the base composition of the position preceding read starts on the horse reference genome showed an excess of Guanine and, to a lesser extent, of Adenine residues (Figure 2—figure supplement 1). This is in line with depurination driving post-mortem DNA fragmentation (Briggs et al., 2007). Additionally, error rate estimates for each nucleotide substitution class indicated the predominance of CàT and GàA mis-incorporations (Figure 2—figure supplement 2). Such mis-incorporation rates were particularly inflated towards read ends, but not read starts (Figure 2—figure supplement 1). This is in line with the DNA nucleotide mis-incorporation profiles expected for the type of DNA library constructed (Seguin-Orlando et al., 2015), and indicates Cytosine deamination at 5’-overhanging ends as the most prominent post-mortem DNA degradation reactions (Jonsson et al., 2013). GATK HaplotypeCaller was used to obtain individual gvcf files with “--minPruning 1 --minDanglingBranchLength 1” to increase sensitivity. Then we employed GATK GenotypeGVCFs for genotyping with the option “--includeNonVariantSites” in order to retain non-variant loci. The vcf files were further filtered in TreeMix and G-PhoCS analysis.

### Principal component analysis (PCA)

The genotype likelhood framework implemented in ANGSD helped mitigate various error rates in ancient and modern genomes. Using EquCab3 (Kalbfleisch et al., 2018) as the reference genome, ANGSD was run using the following options: “-only_proper_pairs 1 -uniqueOnly 1 -remove_bads 1 -minQ 20 -minMapQ 25 -C 50 -baq 1 -skipTriallelic 1 -GL 2 -SNP_pval 1e-6 -rmTrans 1”. This provided a dataset consisting of a total of 16,293,825 transversions when the horse was included, and 10,094,431 transversions when the horse was excluded (i.e. when analyses were restricted to non-caballine genomes only). In these analyses, only specimens sequenced to an average depth of coverage ≥ 1× were retained. Principal Component Analyses were carried out using the PCAngsd package (Meisner & Albrechtsen, 2018) (Figure 2A). To assess the impact of potential reference bias, all analyses were repeated after mapping the sequence data against the donkey reference (Figure 2—figure supplement 4).

### Phylogenetic inference

#### Mitochondrial phylogeny

Cleaned reads were mapped against the mitochondrial genome (Genbank accession no. NC_001640), following the same procedure as when mapping against the nuclear genome. Samples showing an average depth-of-coverage <1× were disregarded, leaving a total of 17 individuals for further analyses. After removing duplicates, consensus mitochondrial sequences were generated using ANGSD (-doFasta 2 -doCounts 1 -setMinDepth 3 -uniqueOnly 1 -remove_bads 1 -minQ 25 -minMapQ 25). Multiple alignment was performed together with the comparative mtDNA sequences downloaded from GenBank (Table S4) using MUSCLE v3.8.31 (Edgar, 2004), with default parameters. The alignments were then split into six partitions (1^st^, 2^nd^ and 3^rd^ codon positions, rRNA, tRNA, and control region).

Two Maximum Likelihood (ML) trees based on all 6 partitions and excluding the control region (positions 15,469-16,660 of the horse reference mitochondrial genome) were both reconstructed using RAxML-NG v.0.9.0 (A. M. Kozlov, Darriba, Flouri, Morel, & Stamatakis, 2019) with GTR+GAMMA substitution model. A total of 1,000 bootstrap pseudo-replicates were carried out to assess node robustness (Figure 2—figure supplement 5). BEAST 2.5.1.0 (Bouckaert et al., 2019) was used to perform Bayesian phylogenetic reconstruction and to estimate split times. The six partitions described above were used, for which the best substitution model was determined using modelgenerator (version 0.85, (Keane, Creevey, Pentony, Naughton, & McLnerney, 2006)) and a Bayesian Information Criterion. We applied together with the Coalescent Constant Population model and a strict LogNormal correlated molecular clock for 50 million generations (sampling frequency = 1 every 1,000). Convergence was assessed visually using Tracer v1.6 and posterior date estimates were retrieved using 10% as burn-in. The final consensus tree was produced using TreeAnnotator 2.5.1.0 (Drummond & Rambaut, 2007) and plotted using ITOL (Letunic & Bork, 2016) (Figure 2—figure supplement 6).

### Autosomal phylogeny

As for autosomes, we reconstructed a (ML) phylogenetic tree as implemented in the PALEOMIX phylo pipeline for phylogenomic reconstructions (Schubert, Ermini, et al., 2014). This analysis was based on the coding sequence (CDS) of protein-coding genes annotated in EquCab3.0, partitioning data according to 1^st^, 2^nd^ and 3^rd^ codon positions. Maximum Likelihood phylogenetic inference was performed using ExaML v3.0.21 (Alexey M. Kozlov, Aberer, & Stamatakis, 2015) and RAxML v8.2.12 (Stamatakis, 2014), under the GAMMA substitution model with 100 bootstrap pseudo-replicates (Figure 2C and Figure 2—figure supplement 7A). We also repeated the same procedure after mapping against the donkey reference genome and got the same topology (Figure 2—figure supplement 7B).

Additionally, we extracted biallelic single nucleotide polymorphisms (SNPs) from the dataset generated in Section 5 using bcftools v1.9 (Li et al., 2009). Both variant datasets obtained following mapping against the horse and donkey reference genomes were used in this analysis to rule out reference bias. We applied filters composed of minimum phred-scaled quality score quality (QUAL) = 20, sites for all individuals below 2 or twice the mean coverage, and allowed up to three individuals with missing data per site. After disregarding transitions, a total of 18,803,101 (mapping against horse genome) and 19,459,070 (mapping against donkey genome) transversions were finally used as input for TreeMix (Pickrell & Pritchard, 2012) with parameters “-k 500-root TWI”, and considering an increasing number of migrations edges (0≤m≤3; Figure 2—figure supplements 8 and 9, Table S5).

### Admixture analyses with *D*-statistics

*D*-statistics were calculated to investigate potential introgression between *E. ovodovi* and other non-caballines (Figure 3—figure supplement 1) using the doAbbababa2 programme in ANGSD (Soraggi et al., 2018). Individuals were grouped by their respective species. D-statistics were computed in the form (((H1, H2), H3), Outgroup) considering only the autosomal sites from bam files mapping against the horse reference with the following options: “-minQ 20 -minMapQ 25 -remove_bads 1 -only_proper_pairs 0 -uniqueOnly 1 -baq 1 -C 50”. The horse reference genome was used as the Outgroup. H1 and H2 denoted any non-caballine genomes except *E. ovodovi* while H3 denoted the *E. ovodovi*. Confidence intervals were estimated applying a jackknife procedure and 5-Mb windows. Z-scores with absolute values higher than 3 were considered to be statistically significant. To eliminate the bias of the reference genome, we also rerun the same analysis using sequence alignments against the donkey reference genome (Figure 3—figure supplement 2).

### G-PhoCS demographic model

#### Data preparation and filtering

In order to model the equine evolutionary history, we selected a total of 10 individuals representing each individual lineage and used their high-coverage genomes as input for G-PhoCS (Gronau et al., 2011). Genotypes were called by GATK and candidate ‘neutral’ loci were identified by applying the following filters:

1. The simple repeats track available for the reference genome was obtained from Ensembl v99 release; corresponding regions were masked.
2. All exons of protein-coding genes were discarded together with their 10 kb flanking regions; this was done based on the GTF format annotation file of the reference genome available from Ensembl v99 Genome Browser.
3. We identified conserved noncoding elements (CNEs) using phastCons scores (based on the 20-way Conservation track provided for the mammal clade according to the genomic coordinates of the human reference) downloaded from the Table Browser of UCSC. All CNEs and their 100 bp flanking regions were masked, using liftOver to convert human genome coordinates into EquCab3.0 horse genome coordinates.
4. Exons of noncoding RNA genes together with their one kilobase flanking regions were removed, based on the annotations available for the reference genome.
5. Gaps in the reference genome were disregarded.

Besides the various hard filters described above, regions/sites likely to (1) be enriched for misaligned bases, and to (2) have high false negative rates during read alignment or variant detection were masked as missing data. Different individuals may be treated differently depending on the result of genotyping in Section 5 depending on the presence of (1) indels, (2) triallelic sites, (3) positions with depth of coverage twice the mean depth recorded for each individual, and; (4) transition sites.

We selected 1 kb loci located with minimum inter-locus distance of 30 kb from the intervals that pass all the criteria described above. Then consensus sequences were generated for each individual from the vcf file generated in Section 5 using bcftools ‘consensus’ command, with IUPAC codes indicating heterozygous genotypes (--iupac-codes) and “N” representing masked sites (--mask and --missing ‘N’).

Finally, we excluded contiguous intervals if the total amount of missing bases was greater than 50% of the region length, resulting in a final collection of 15,324 loci using the horse reference genome (autosomes only). Neighbor-joining trees were constructed to confirm the topology before the inferring the population divergence (Figure 3—figure supplement 3).

### MCMC setup

We used default global settings (Gronau et al., 2011), including a Gamma prior distribution (α= 1, β= 10,000) for all mutation-scaled population sizes (*θ*) and a Gamma prior distribution (α= 0.002, β= 0.00001) for all mutation-scaled migration rate (*m*). The initial parameter value of mutation-scaled divergence times (*τ*) was first set individually for each population. Then we ran ∼100,000-200,000 iteration tests and manually evaluated the convergence by checking the achieve acceptance ratios (*ie*. accept if around 30-70%) or using Tracer v1.6 (http://tree.bio.ed.ac.uk/software/tracer/). For each test, we updated the input of the initial *τ* and all fine-tuned parameters based on previous results to get the appropriate value. The final results in Figure 3 are based on 500,000 MCMC iterations.

### Parameter calibration

We assumed an average generation time (*g*) of 8 years, considering the mutation rate *μ* (per year) could be variable when using different sequences. The coalescent time of the *Equus* (4.0-4.5 Mya) (L. Orlando et al., 2013) was used to bound the parameter *μ*. Effective population sizes (*Ne*) and divergence times (*T*) were estimated by calibrating *θ* and *τ* parameter using *g* and *μ* (Table S6), given by: *Ne* = *θ*/(4*μg*) and *T* =*τ*/*μ* (Gronau et al., 2011).

### Inferring gene flow

Total migration rates (*M*) were estimated by a mutation-scaled version (*m*) given by: *M* = *mτ_m_*, where *τ_m_* is the mutation-scaled time span of the migration band. We then converted such rates, *M* , into a probability of migration using the formula: *p* = 1-e^−*M*^ (where *p* is the probability of gene flow), according to the method presented in (vonHoldt et al., 2016).

The migration model implemented in G-PhoCS makes it possible to detect gene flow between any two lineages by introducing migration bands manually to the demographic model. However, it remains difficult to detect weak migration events. Additionally, scenarios including a large number of migration bands can lead to spurious results. To address this, we first inferred a demographic model with no migration bands, and then introduced several migration bands corresponding to five independent scenarios (Table S6). A significant migration band was considered supported if both the 95% Bayesian credible interval of total migration rate (*M*) did not include 0 and the mean value of *M* was estimated to be greater than 0.03.

Settings for the migration bands between extant caballines are based on previous research (Jonsson et al., 2014). The significant migration band from horse to the non-caballine ancestor were identified (Table S6), in line with previous work (Jonsson et al., 2014). However, no other non-negligible (*M* > 3%) migration bands was found in our analyses (Table S6).

We then tried to estimate the migration events between *E. ovodovi* and other branches. We added all possible migration bands between *E. ovodovi* and non-caballine branches into the demographic model except the migration bands between *E. ovodovi* and the ancestor of non-caballines, as the model is often underpowered to infer migration between sister populations. All of the migration bands were separated into four demographic models. Only three migration bands were shown significant (Table S6).

Finally, the total four migration bands were combined into one demographic model (Table S7) and compared the estimates to the one including no migration (Table S8).

### Demographic trajectories with PSMC

#### PSMC analyses

In order to reconstruct the past demographic dynamics of the *E. ovodovi* lineage, we applied the PSMC algorithm (version 0.6.5-r67) (Li & Durbin, 2011) to the sample HH06D (12.0×) , as well as three other Eurasian equine species (*E. caballus* TWI, *E. hemionus* ONA and *E. kiang* KIA).

We first obtained the diploid consensus sequences after mapping against the horse genome for the autosomes of each specimens using bcftools ‘mpileup’ command and the ‘vcf2fq’ command from vcfutils.pl with the following filters: mapping quality ≥ 25; adjust mapping quality =50; minimum depth-of-coverage = 8; maximum depth-of-coverage ≤ 99.5% quantile of the coverage distribution; minimum RMS mapping quality = 10; filtering window size of indels = 5.

After filtering the bases with Phred quality scores strictly lower than 35, we ran PSMC with following command: ‘psmc -N25 -t15 -r5 -p “4+25*2+4+6” ’. Calibration was carried out using a generation of 8 years and mutation rate of 7.242×10^−9^ per generation per site, following previous work (Jonsson et al., 2014). However, as for the mis-incorporation pattern and high error rate of HH06D (Figure 2—figure supplements 1 and 2), we also performed analyses without transitions using mutation rates of 2.3728×10^−9^ that was obtained assuming that the most recent common ancestor of living equine species emerged 4 Mya (L. Orlando et al., 2013).

We found a great expansion of HH06D in the past 50,000 years when retaining transitions but not when conditioning on transversions (Figure 4—figure supplement 1). The former is thus likely spurious and at least partly driven by severe post-mortem DNA damage signatures in the sequence data. We therefore only used the latter when considering the ancient HH06D specimen.

#### False negative rate correction

The HH06D genome (12.0×) was corrected assuming a uniform false-negative rate (uFNR) following (L. Orlando et al., 2013), as the average depth-of-coverage is lower than the recommended 20×. To identify the correction value of uFNR for HH06D, we randomly down-sampled reads of SOM genome (21.0×) using DownsampleSam function of Picard Tools to down-scale sequence data to the same average depth-of-coverage as that obtained for HH06D. This indicated that a value of 0.22 was the most suitable uFNR value for rescaling the HH06D PSMC profile (Figure 4—figure supplement 2A). The KIA and the ONA genomes, which also showed limited coverage, were also rescaled following the same procedure (Figure 4—figure supplement 2B-C). Finally, PSMC confidence intervals were assessed from 100 bootstrap pseudo-replicates (Figure 4).

### Heterozygosity Inference and Inbreeding

Global heterozygosity rates and inbreeding levels were inferred for high coverage individuals (>10×) using ROHan (Renaud, Hanghoj, Korneliussen, Willerslev, & Orlando, 2019) with default parameters, except that transitions were excluded (--tvonly) (Figure 5—figure supplement 1). To limit the impact of remnant mis-incorporations, we used the attached estimateDamage.pl script to estimate damage for all ancient samples prior to heterozygosity computation. Inbreeding was co-estimated together with genome-wide heterozygosity levels from the total ROH length (Figure 5—figure supplement 2).

---

**Figure 1—figure supplement 1.**
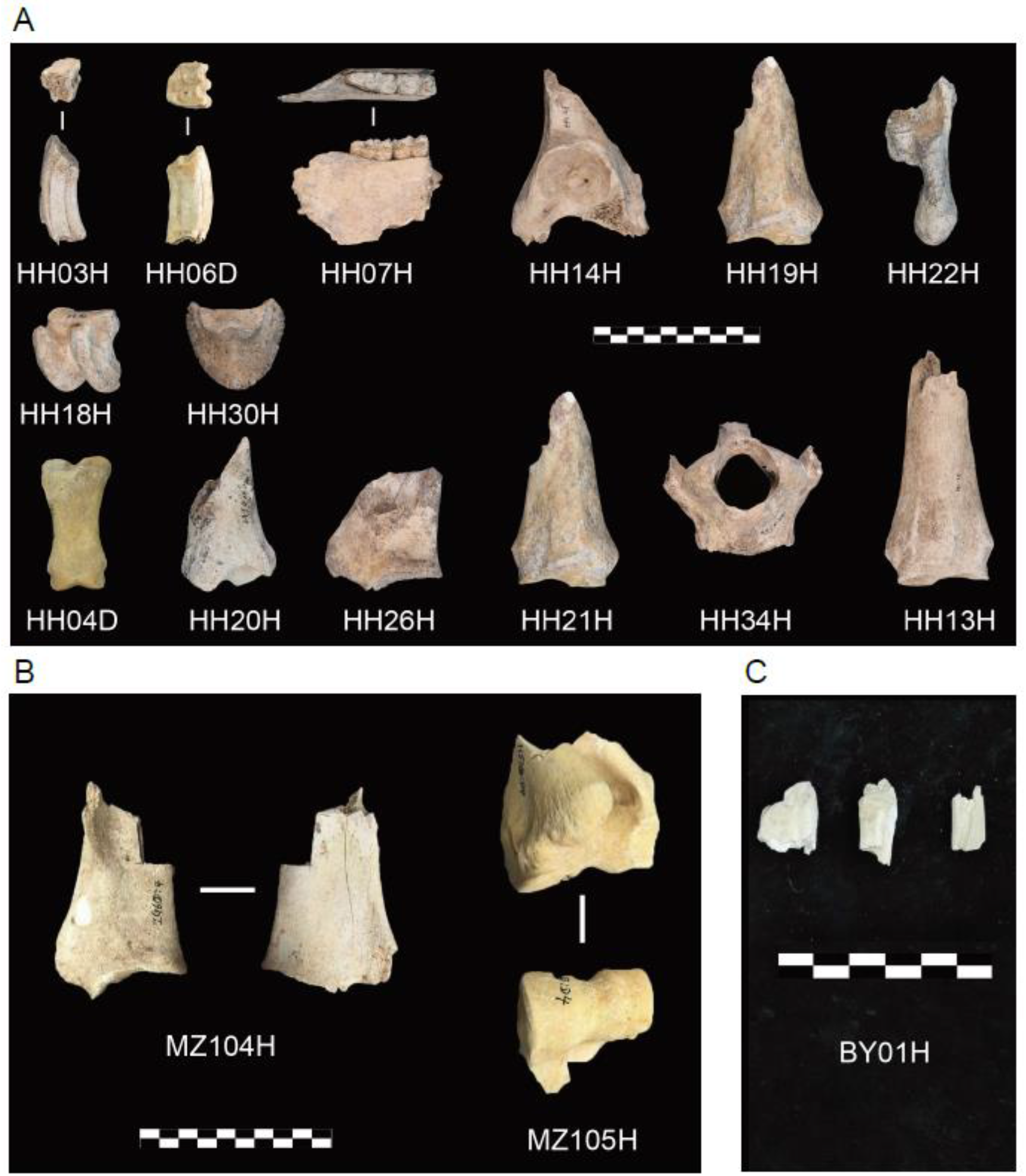
Archaeological material investigated in this study. (A) Honghe (HH), (B) Muzhuzhuliang (MZ) and (C) Shatangbeiyuan (BY).

**Figure 1—figure supplement 2.**
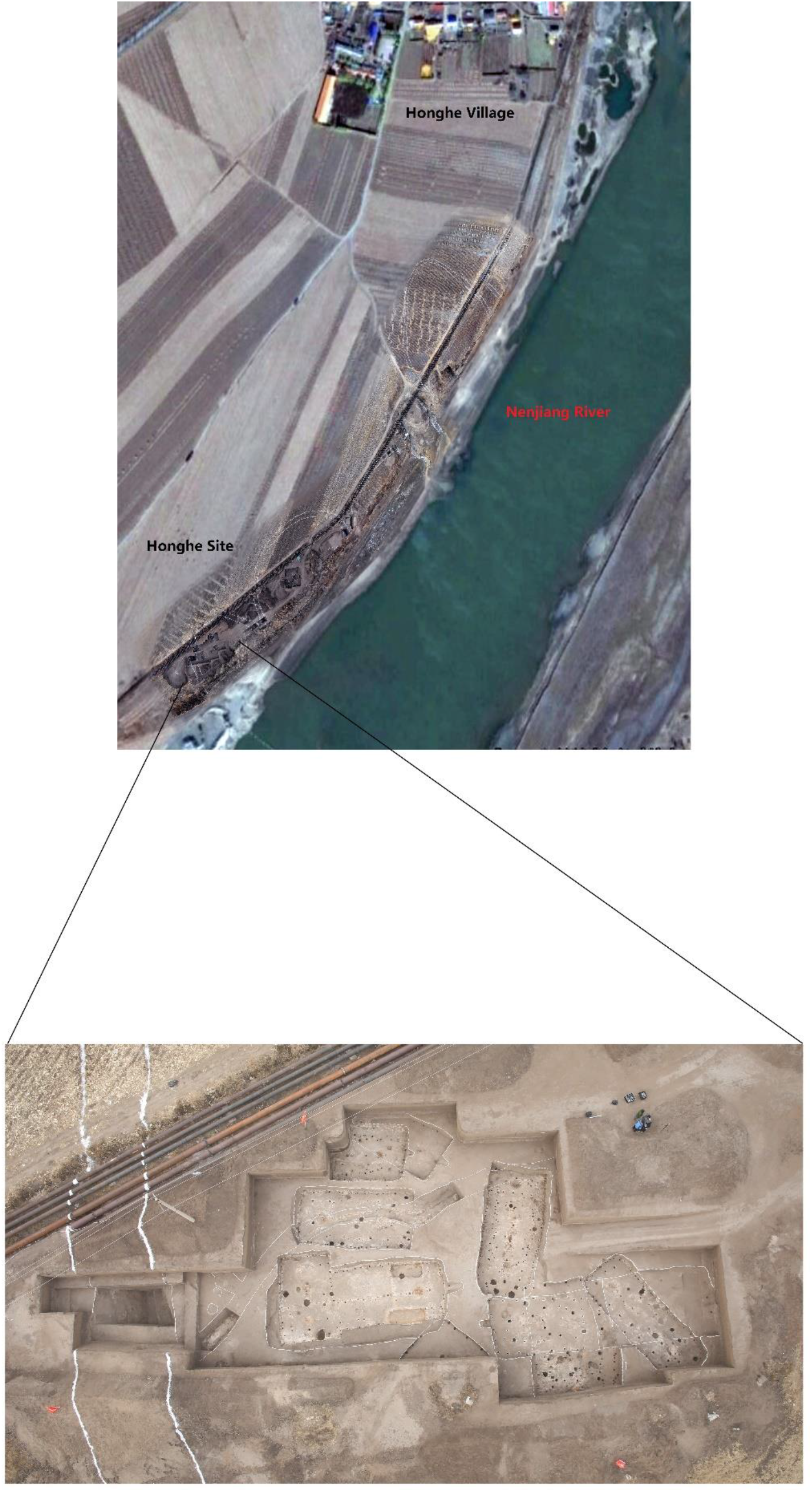
Aerial view of the Honghe site.

**Figure 2—figure supplement 1.**
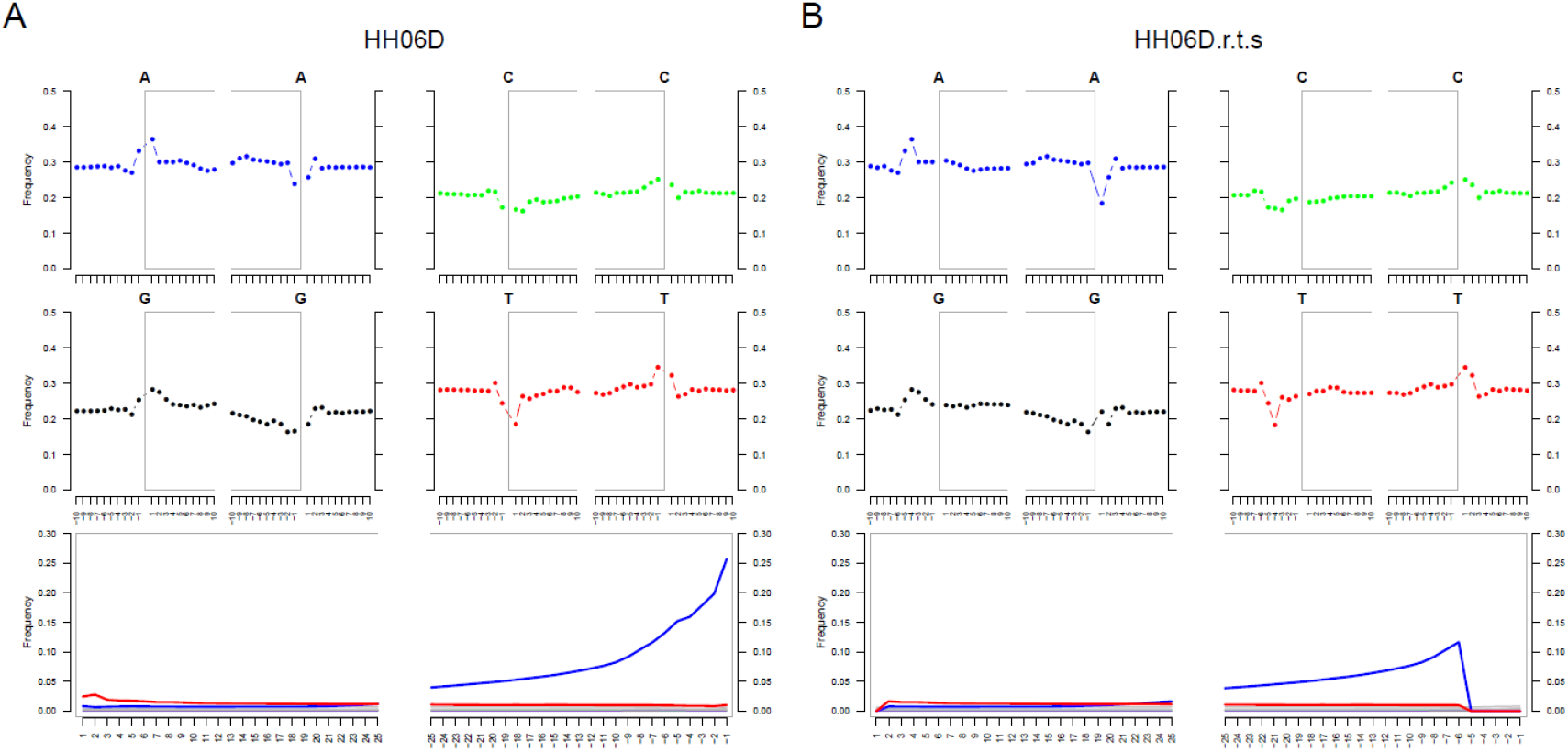
DNA Damage patterns for HH06D. (**A**) Before rescaling and trimming and (**B**) after rescaling and trimming the region comprising the five first and last nucleotides sequenced.

**Figure 2—figure supplement 2.**
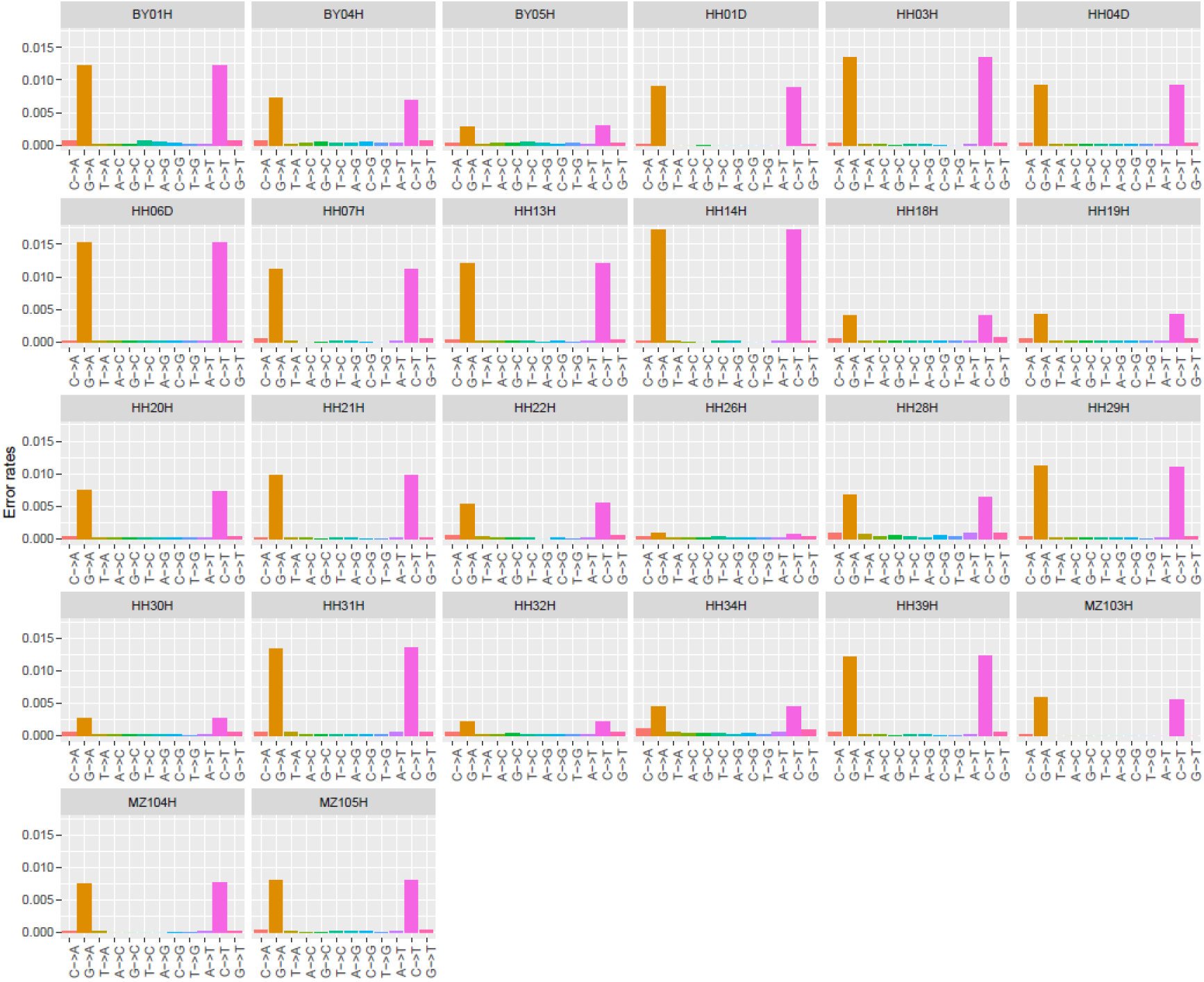
Error profiles of the 26 ancient genomes characterized in this study. After trimming and rescaling, reads showing mapping quality scores inferior to 25 and bases showing quality scores inferior to 20 were disregarded.

**Figure 2—figure supplement 3.**
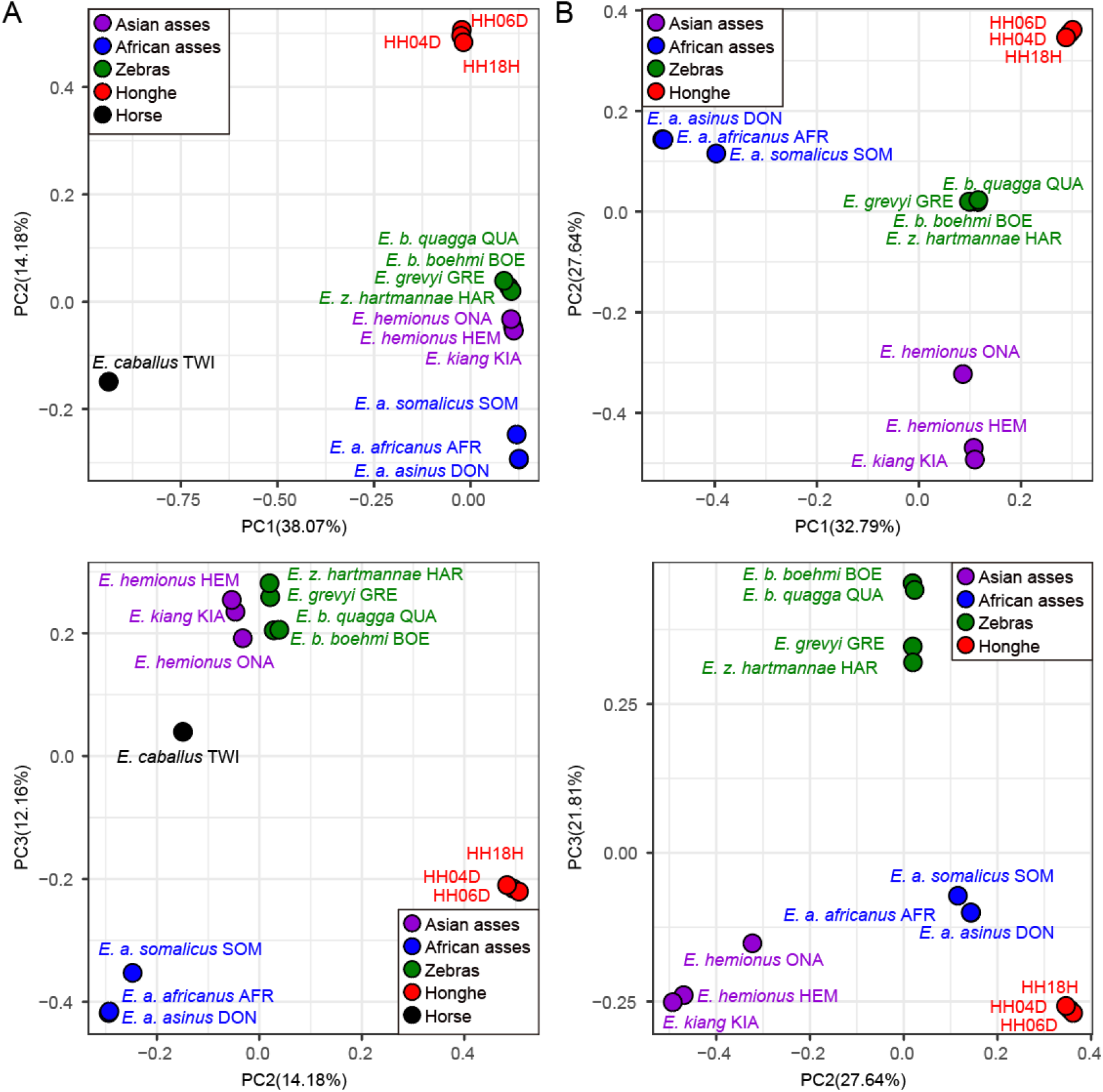
Principal Component Analysis (PCA) based on genotype likelihoods using the horse reference genome. (**A**) Including and (**B**) excluding the outgroup individual underlying the horse reference genome (TWI) (Kalbfleisch et al., 2018). Sequence data were aligned against the horse reference genome (Kalbfleisch et al., 2018).

**Figure 2—figure supplement 4.**
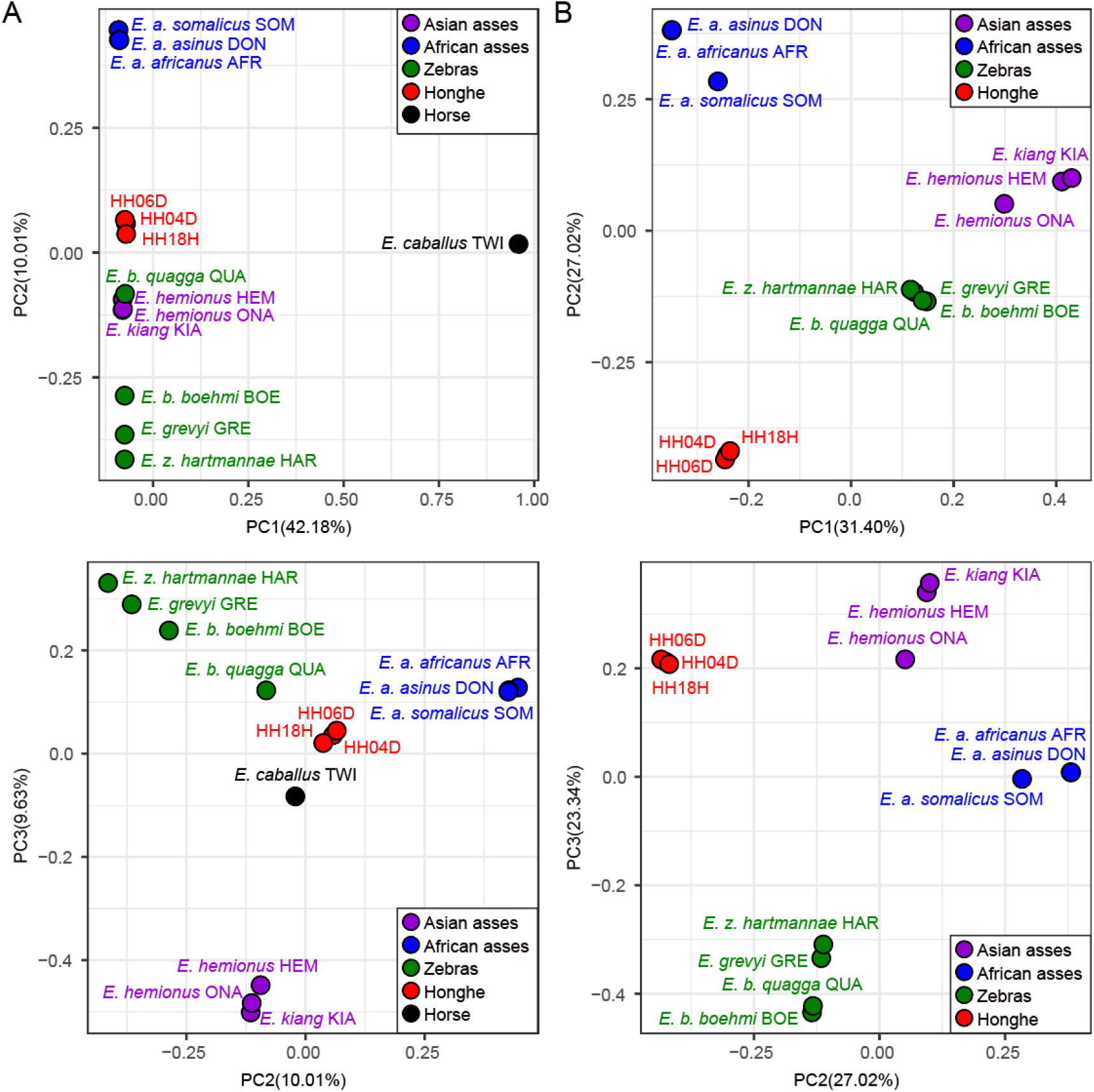
Principal Component Analysis (PCA) based on genotype likelihoods using the donkey reference genome. (**A**) Including and (**B**) excluding the outgroup individual underlying the horse reference genome (TWI) (Kalbfleisch et al., 2018). Sequence data were aligned against the donkey reference genome (Renaud et al., 2018).

**Figure 2—figure supplement 5.**
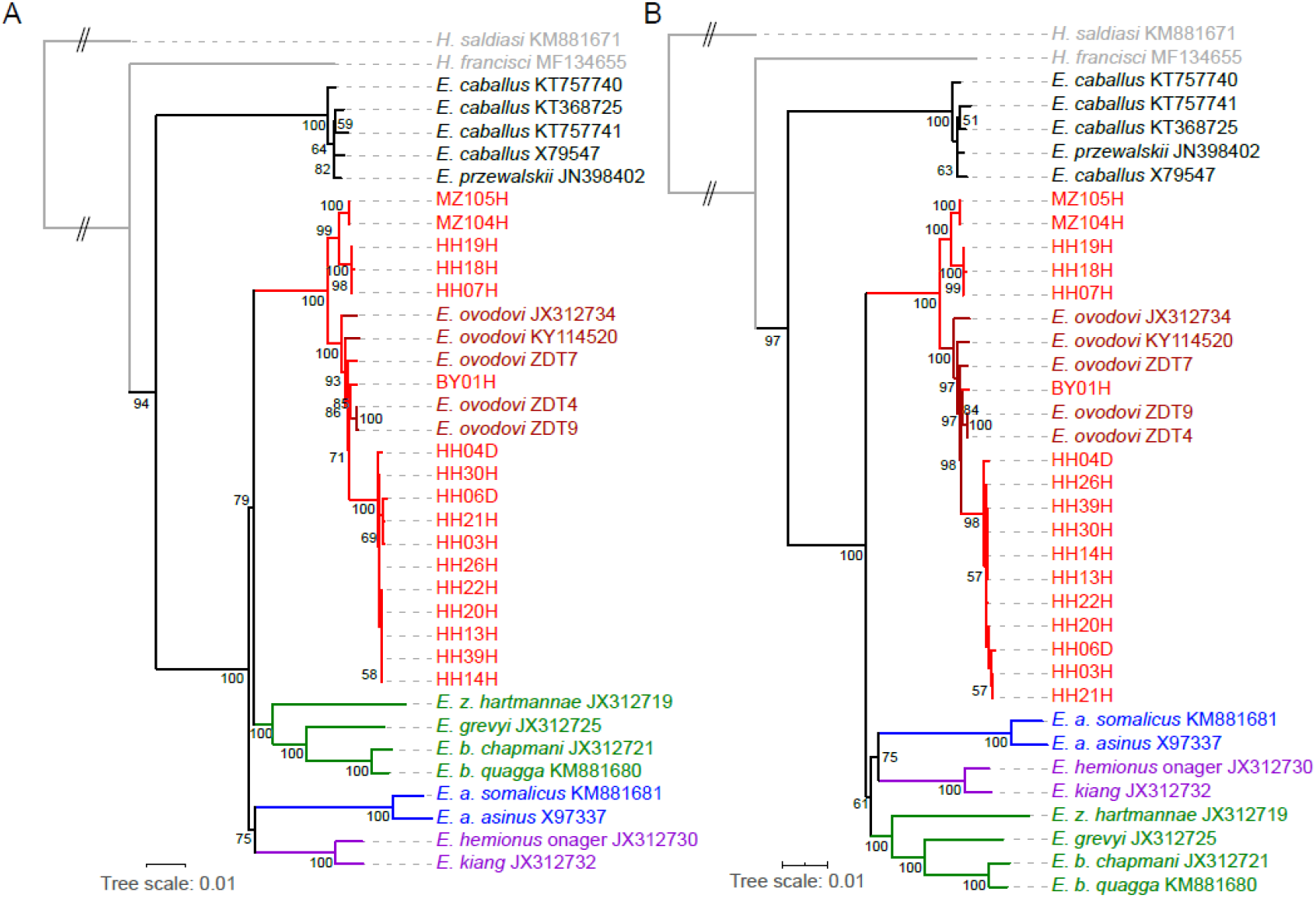
RAxML-NG (GTR+GAMMA model) Maximum Likelihood phylogeny of complete mitochondrial sequence data. (**A**) Including the control region. (**B**) Excluding the control region. Node support was estimated from 1,000 bootstrap pseudo-replicates and the tree was manually rooted using *Hippidion Saldiasi*.

**Figure 2—figure supplement 6.**
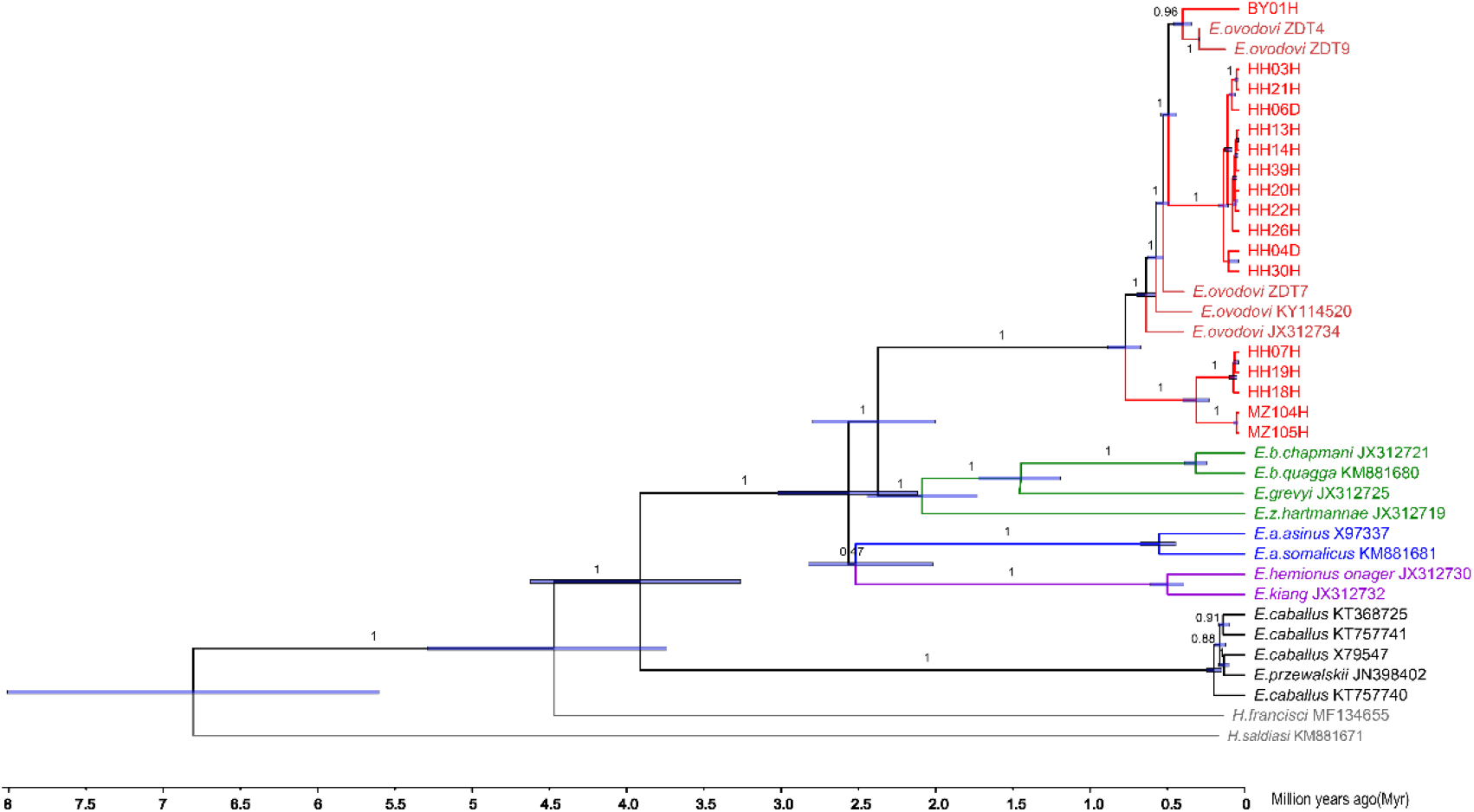
Bayesian mitochondrial phylogeny based on 6 partitions and using *Hippidion Saldiasi* as outgroup. The tree was reconstructed using a total number of 50 million MCMC states in BEAST (sampling frequency = 1 every 10,000, burn-in = 10%). The substitution models applied to the six sequence partitions were the TrN+I+G model (1^st^ codon position = 3,802 sites), the TrN+I model (2^nd^ codon position = 3,799 sites), the GTR+I+G model (3^rd^ codon position = 3,799 sites), the HKY+I model (transfer RNAs = 1,517 sites), the TrN+I+G model (ribosomal RNAs = 2,556 sites) and the HKY+I+G model (control region = 1,192 sites).

**Figure 2—figure supplement 7.**
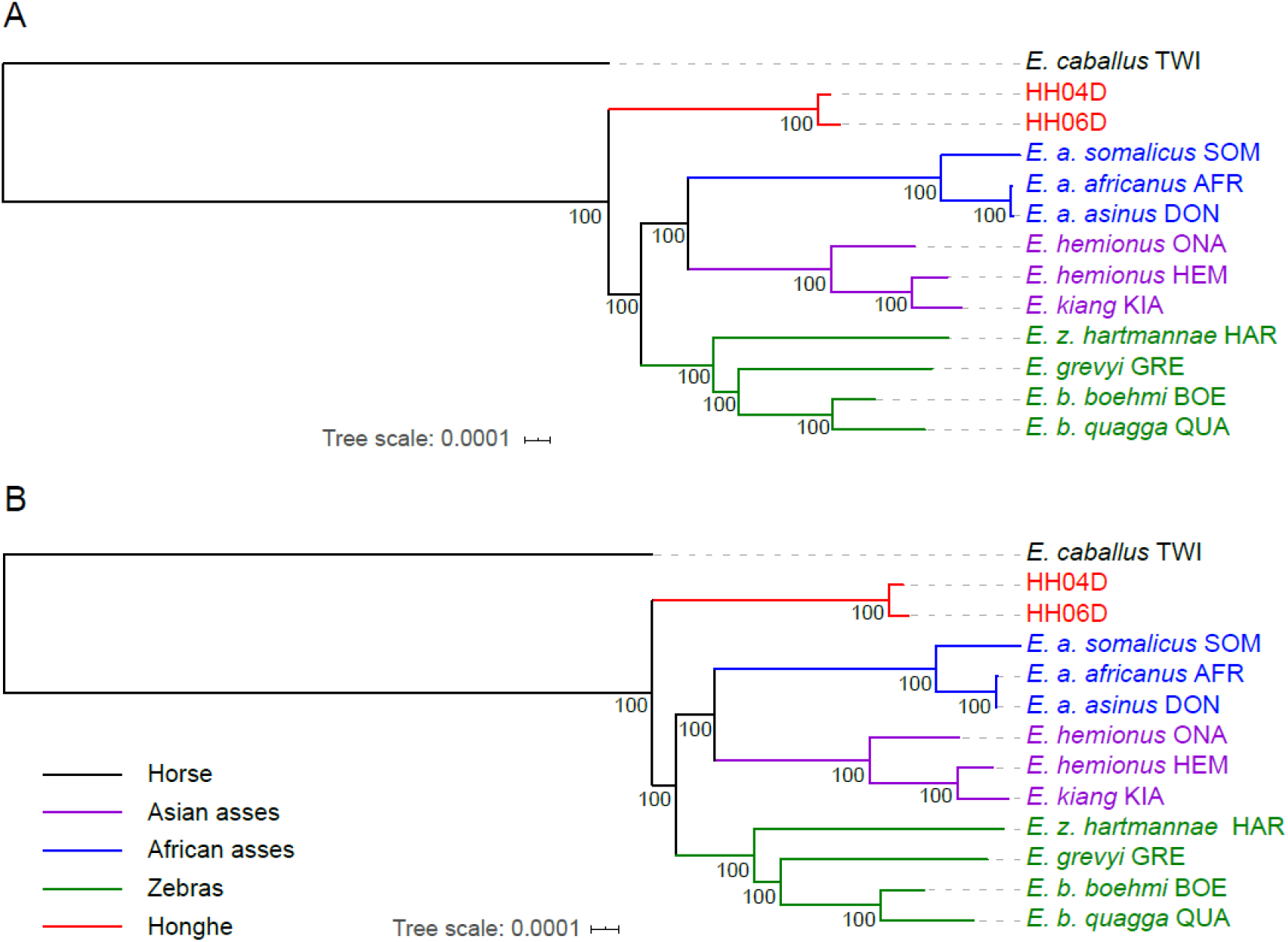
Exome-based Maximum likelihood phylogeny rooted by the horse lineage. (**A**) Using sequence alignments against the horse reference genome (Kalbfleisch et al., 2018). (**B**) Using sequence alignments against the donkey reference genome (Renaud et al., 2018). Node supports were estimated from 100 bootstrap pseudo-replicates.

**Figure 2—figure supplement 8.**
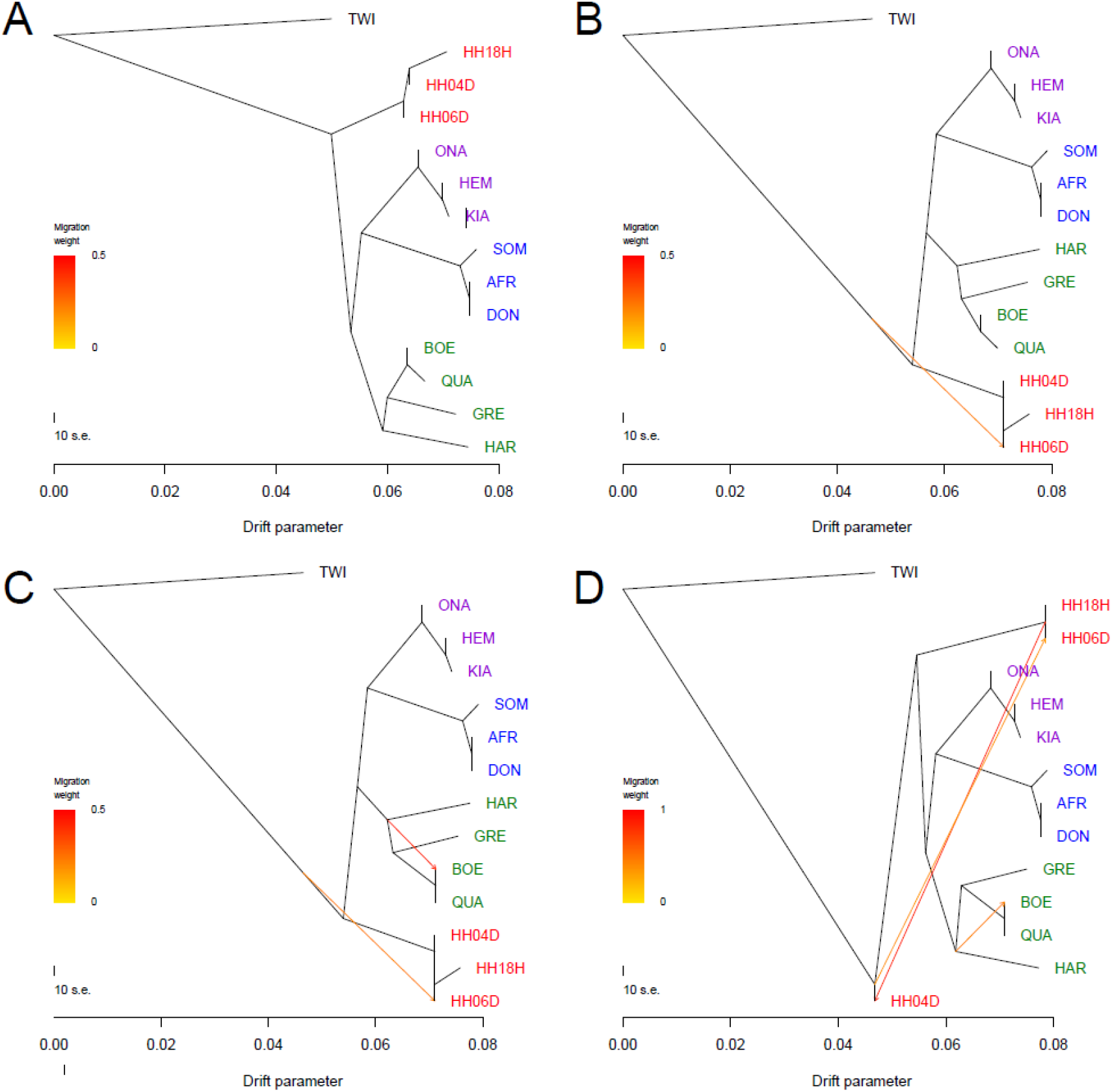
Treemix analysis of based on genome-wide SNP data conditioned on transversions using the horse reference genome. Sequence data were mapped against the horse reference genome (Kalbfleisch et al., 2018). A total of 0 to 3 migration edges were considered. The result of each analysis is shown in panels (**A**) to (**D**), respectively. Considering additional migration edges did not improve the variance explained by the TreeMix model (Table S5).

**Figure 2—figure supplement 9.**
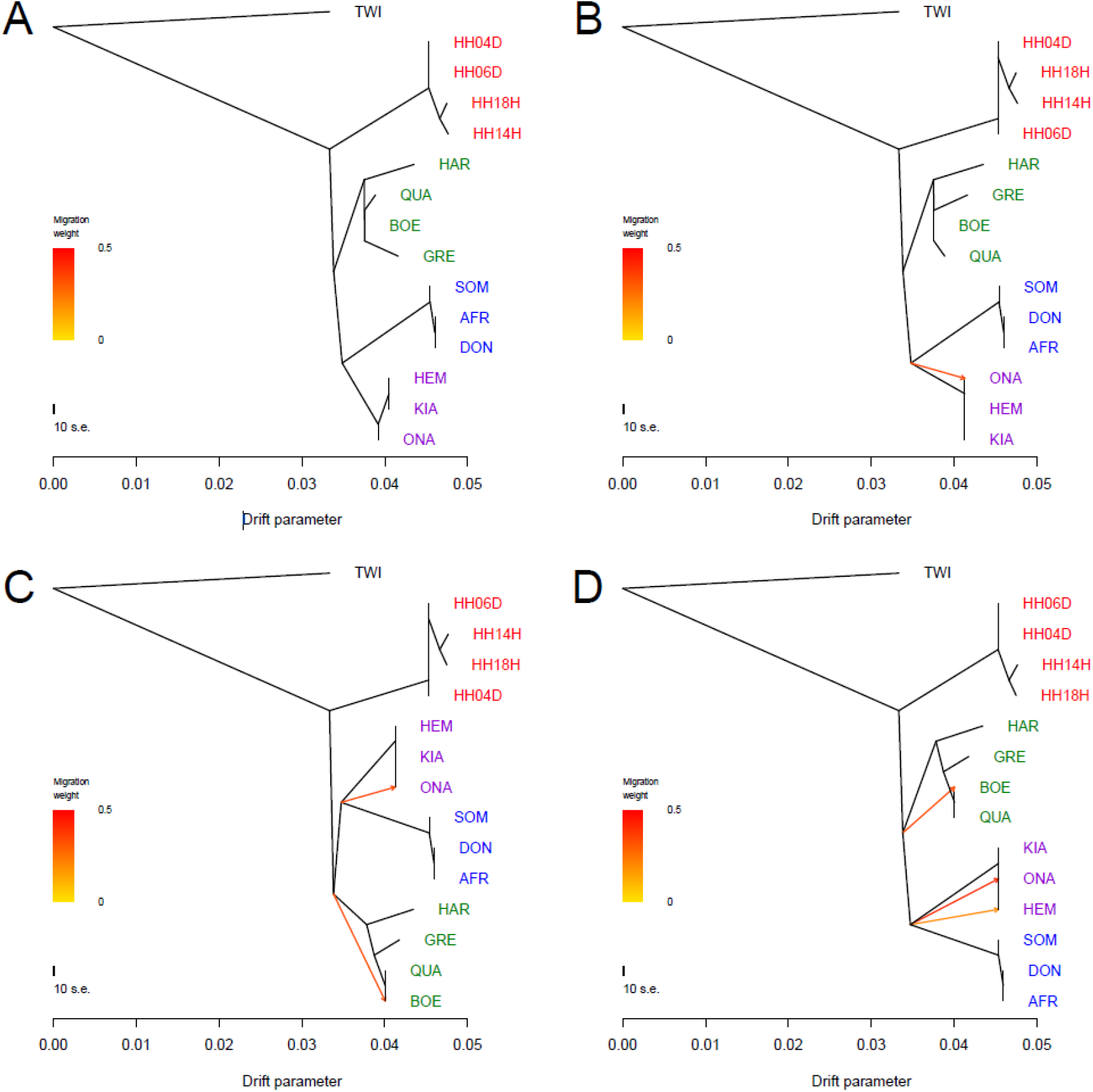
Treemix analysis of based on genome-wide SNP data conditioned on transversions using the donkey reference genome. Sequence data were mapped against the donkey reference genome (Renaud et al., 2018). A total of 0 to 3 migration edges were considered. The result of each analysis is shown in panels (A) to (**D**), respectively. Considering additional migration edges did not improve the variance explained by the TreeMix model (Table S5).

**Figure 3—figure supplement 1.**
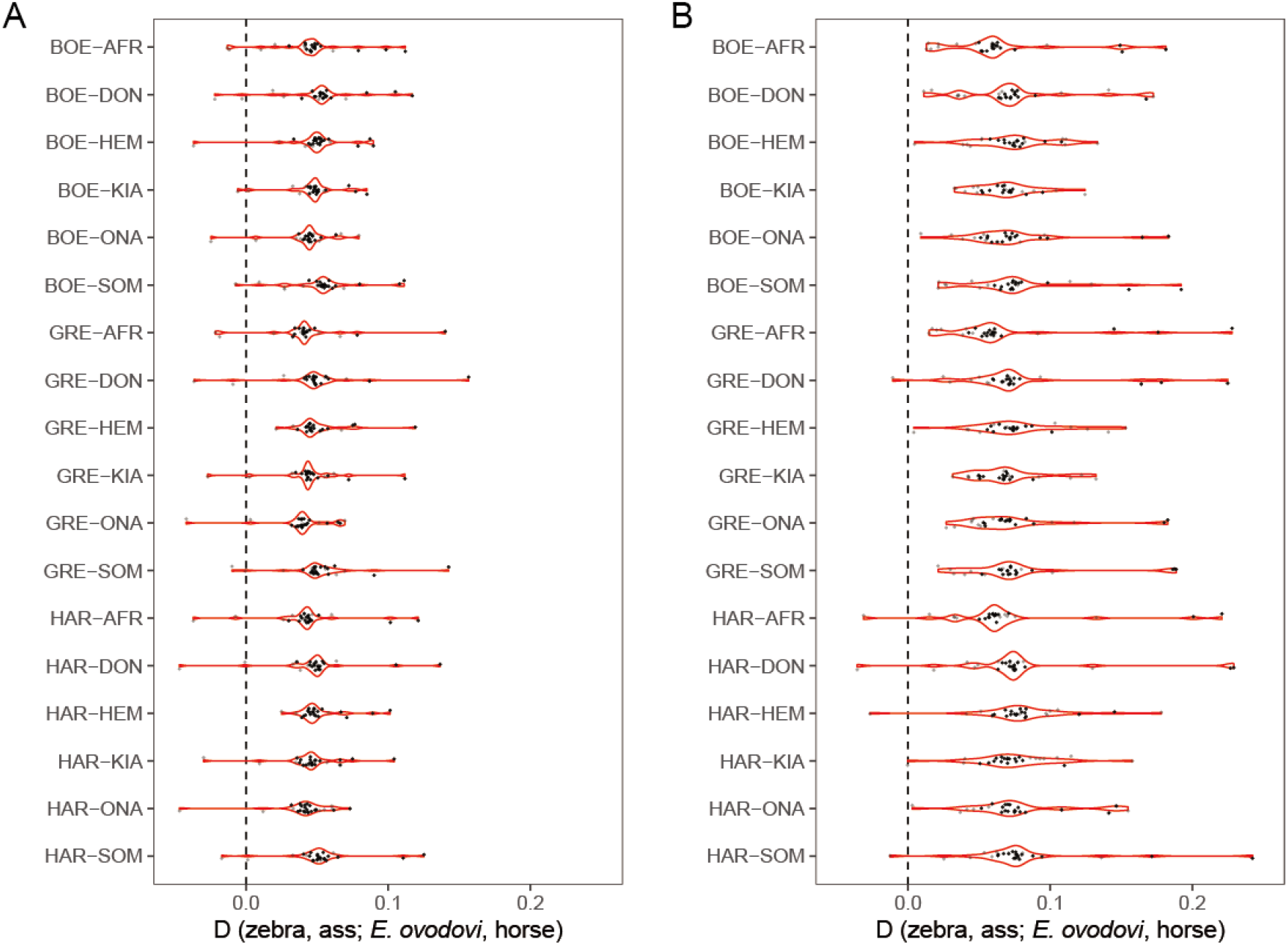
*D-*statistics in the form of (zebra, ass; *E. ovodovi*, outgroup), using sequence alignments against the horse reference genome. Significantly positive D-statistics are indicative of an excess of shared derived polymorphisms between *E.* (*Sussemionus*) *ovodovi* and extant assess, which is compatible with admixture between both lineages. (**A**) Including transitions. (**B**) Excluding transitions.

**Figure 3—figure supplement 2.**
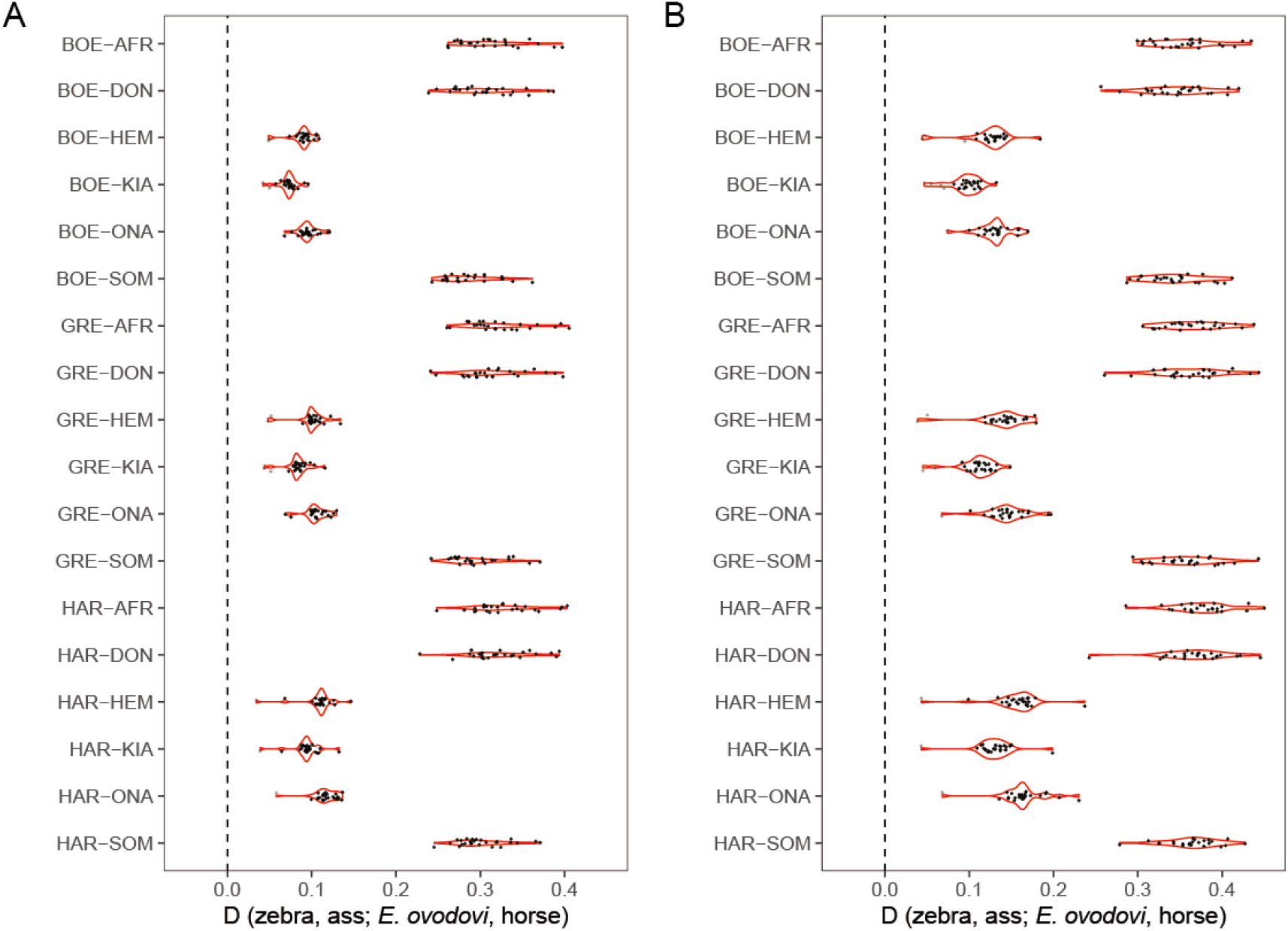
*D-*statistics in the form of (zebra, ass; *E. ovodovi*, outgroup), using sequence alignments against the donkey reference genome. Significantly positive D-statistics are indicative of patterns of shared derived polymorphisms between *E.* (*Sussemionus*) *ovodovi* and extant assess, which is compatible with admixture between both lineages. (**A**) Including transitions. (**B**) Excluding transitions.

**Figure 3—figure supplement 3.**
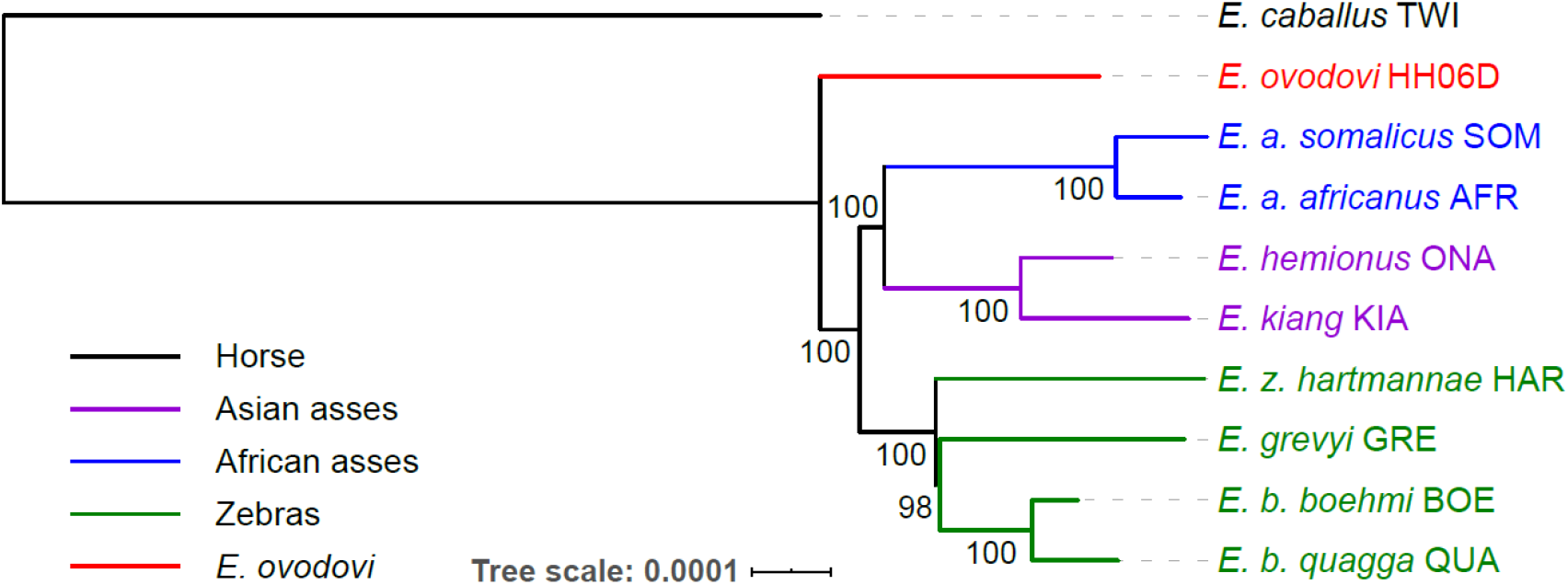
NJ tree of selected samples based on 15,324 candidate ‘neutral’ loci identified using sequence alignments against the horse reference genome (detailed in **Data preparation and filtering**). Node supports were assessed from 1,000 bootstrap pseudo-replicates.

**Figure 4—figure supplement 1.**
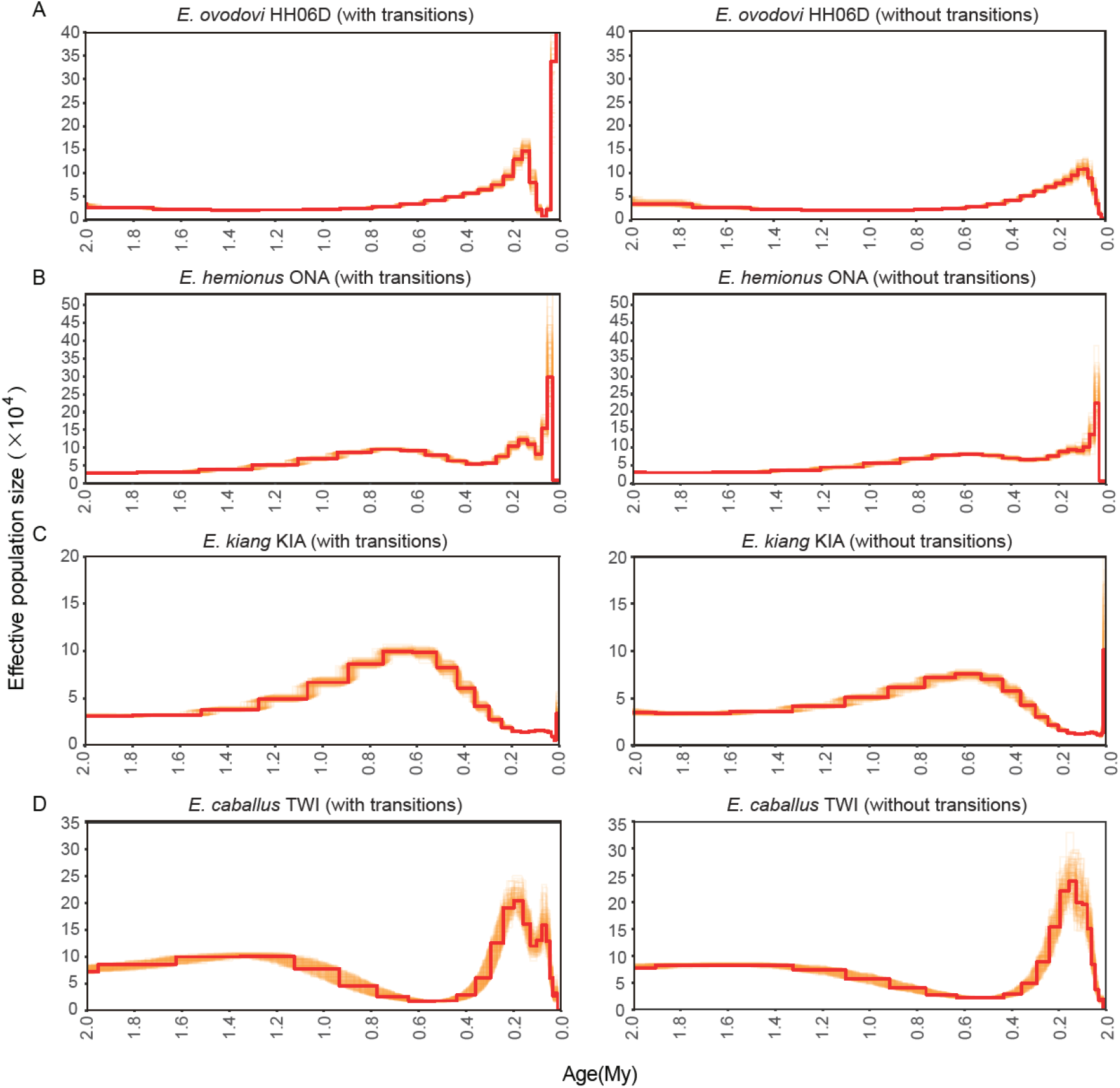
PSMC bootstrap pseudo-replicates for samples with (left) and without (right) transitions. (**A**) HH06D, (**B**) ONA, (**C**) KIA and (**D**) TWI. The *E. ovodovi* genome still included, even after rescaling, a significant proportion of nucleotide mis-incorporations pertaining to post-mortem DNA damage. This resulted in the presence of an excessive fraction of singleton mutations along this lineage, and the artefactual expansion observed in the most recent time range.

**Figure 4—figure supplement 2.**
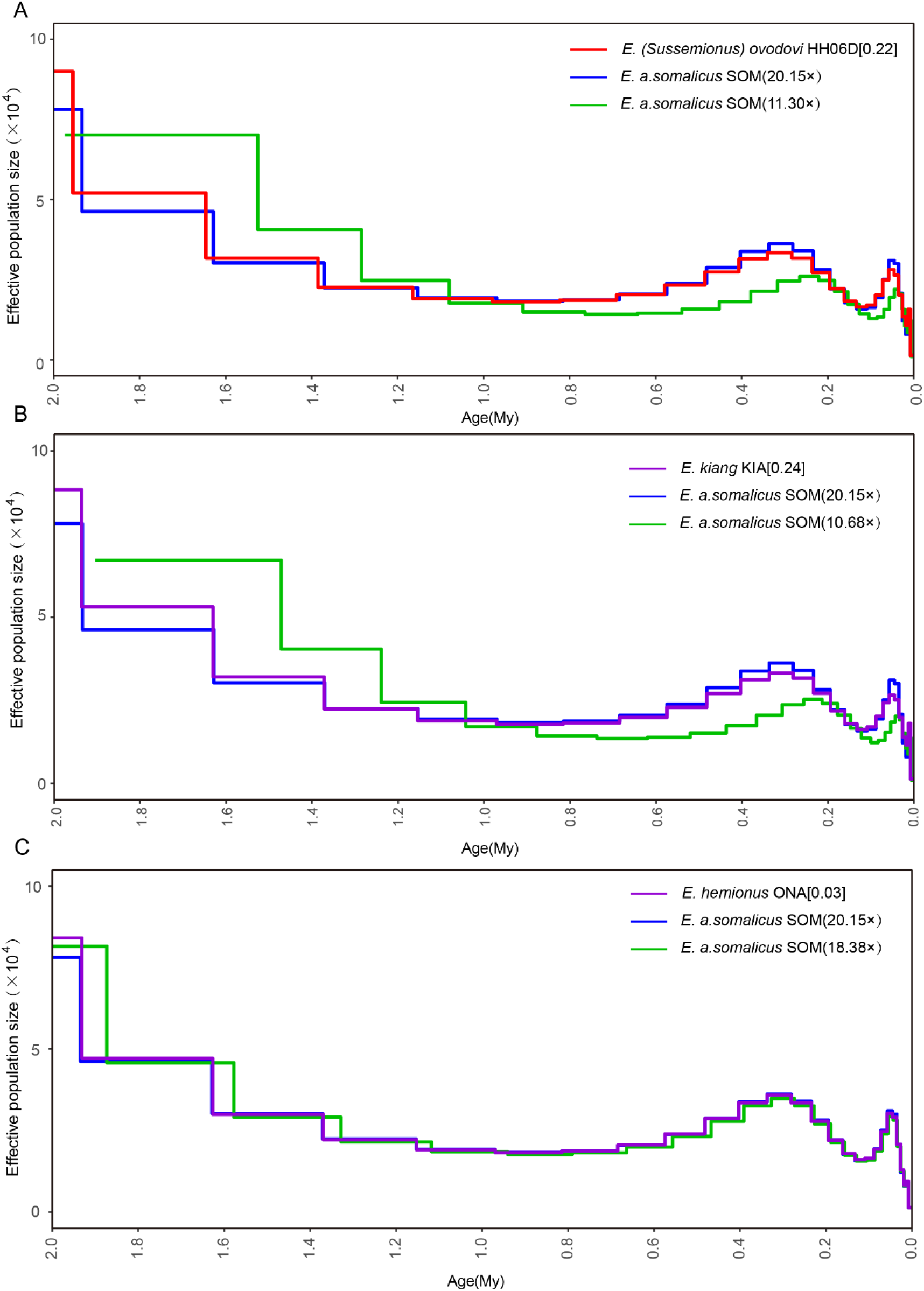
Determining the uniform false-negative rate (uFNR) that was necessary for PSMC scaling. (**A**) HH06D (11.30×), (**B**) KIA (10.68×) and (**C**) ONA (18.38×).

**Figure 5—figure supplement 1.**
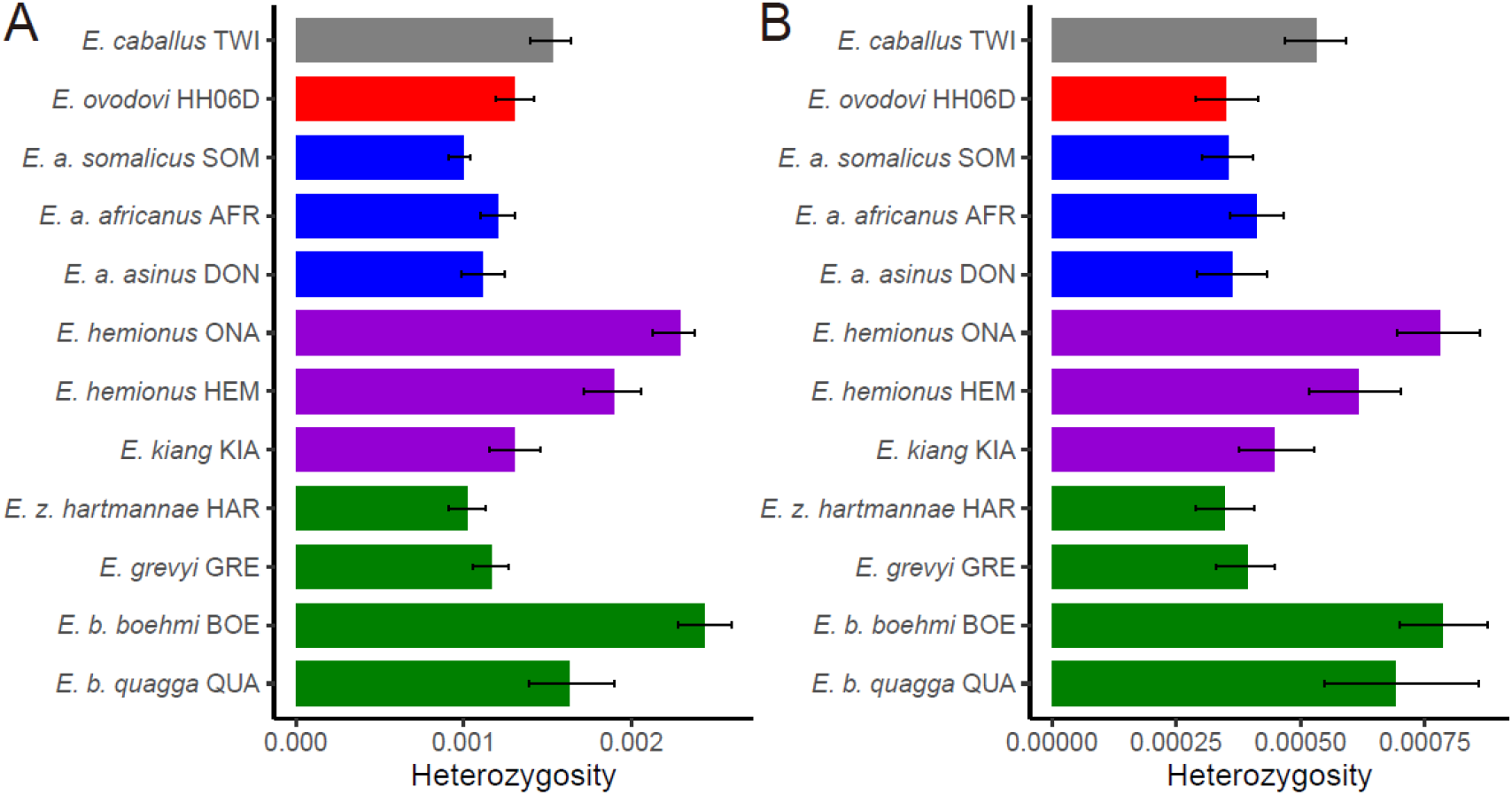
Heterozygosity rates outside Runs-of-Homozygosity (ROH) together with 95% confidence intervals. (**A**) Including transitions and (**B**) excluding transitions.

**Figure 5—figure supplement 2.**
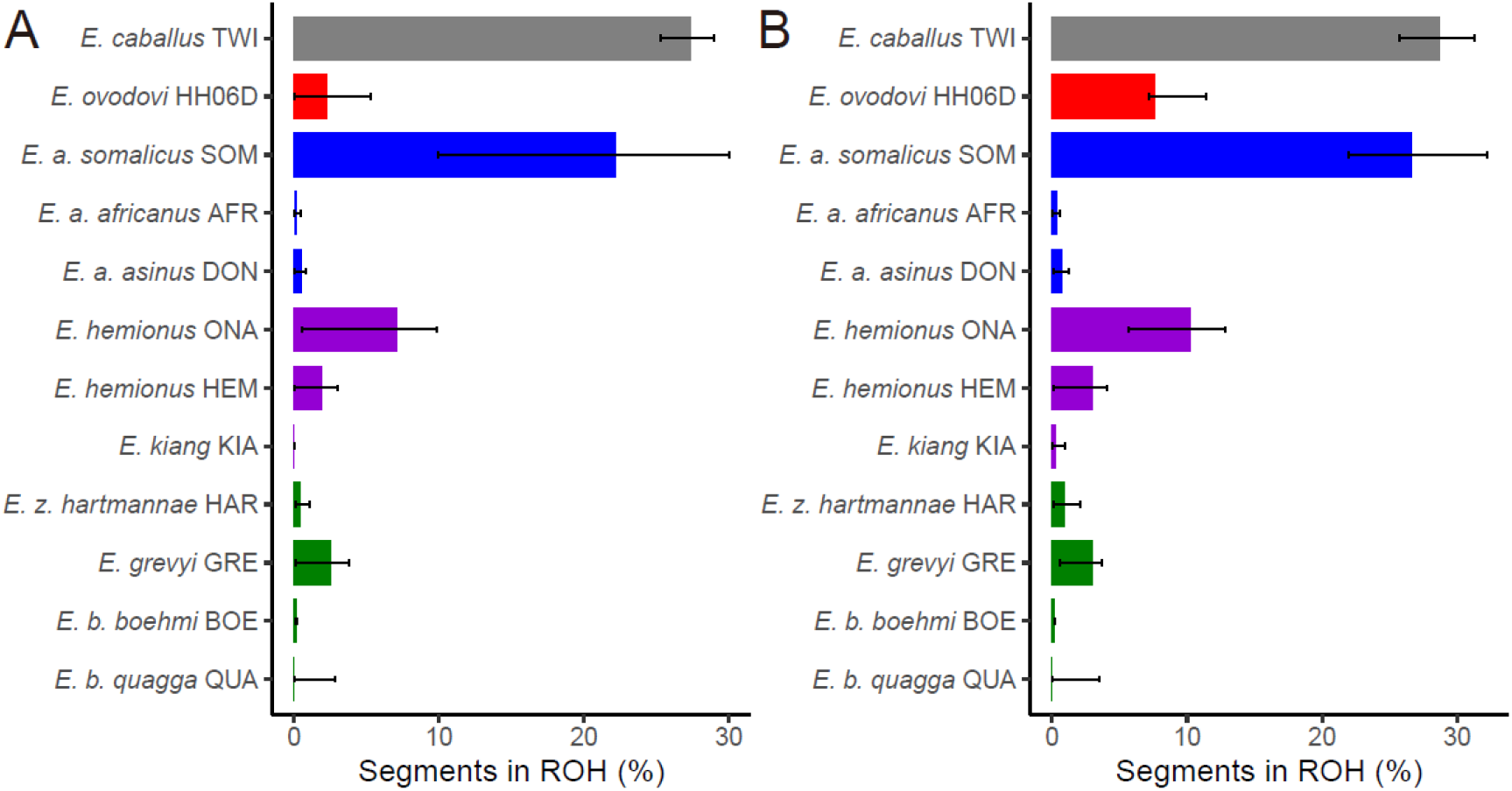
The fraction of the genome segments consisting of ROHs together with 95% confidence intervals. (**A**) Including transitions and (**B**) excluding transitions.

